# Telomerase Prevents Emphysema in Old Mice by Sustaining Subpopulations of Endothelial and AT2 Cells

**DOI:** 10.1101/2021.01.07.425708

**Authors:** Marielle Breau, Christelle Cayrou, Dmitri Churikov, Charles Fouillade, Sandra Curras-Alonso, Serge Bauwens, Frederic Jourquin, Laura Braud, Frederic Fiore, Rémy Castellano, Emmanuelle Josselin, Carlota Sánchez-Ferrer, Giovanna Giovinazzo, Eric Gilson, Ignacio Flores, Arturo Londono-Vallejo, Serge Adnot, Vincent Géli

## Abstract

Accumulation of senescent cells has been causally linked to the development of age-related pathologies. Here, we characterized a new mouse model (p21^+/Tert^) whose telomerase (TERT) is expressed from the p21 promoter that can be activated in response to telomere dysfunction. Lung parenchyma from p21^+/Tert^ old mice accumulated fewer senescent cells with age and this correlated with a reduction in age-related alveolar space enlargement, a feature of pulmonary emphysema. This protection against emphysema depends on TERT catalytic activity and is associated with increased proliferation of pulmonary endothelial cells (EC) and capillary density. Single-cell RNA sequencing of lung cells revealed that TERT expression was associated with the enrichment of ECs expressing genes involved in vessel regeneration and in AT2 cells overexpressing S/G2M markers. These findings indicate that p21-promoter-dependent expression of catalytically active telomerase prevents emphysema by sustaining the proliferation of subclasses of EC and AT2 cells.

## Introduction

There is a growing consensus that accumulation of senescent cells in tissues represents a key process that contributes to age-related health decline ^1,2^. In humans, an increasing number of age-related diseases have been associated with abnormally short telomeres including dyskeratosis congenita, aplastic anaemia, pulmonary fibrosis, and lung emphysema ^3^. Telomere erosion appears as a major risk factor for never smoker patients with pulmonary fibrosis while in smokers mutations that affect telomerase activity favour the development of emphysema/chronic obstructive pulmonary disease (COPD) ^4–7^. In mouse, induction of DNA damage with critically short or dysfunctional telomeres results in the development of pulmonary fibrosis ^8^. In addition, telomerase-deficient mice treated with cigarette smoke were shown to develop pulmonary emphysema due to the release of inflammatory cytokines by senescent cells ^4,9,10^. These results inspired development of adeno-associated vectors (AAV) encoding the telomerase reverse transcriptase gene (TERT) to assess the therapeutic effect of telomerase ectopic expression either in old mice or in mice with experimentally shortened telomeres ^11–13^. Telomerase gene therapy had beneficial effects in delaying physiological aging and improving pulmonary function. However, the precise mechanism by which TERT expression prevents lung damage or promotes lung endogenous repair remains to be elucidated.

The cyclin-dependent kinase inhibitor p21 is the main effector of p53, which is activated in response to DNA damage ^14^ and telomere dysfunction ^15^. The p53-dependent up-regulation of p21 leads to cell cycle arrest in the G1 phase and is the primary event inducing replicative senescence upon telomere shortening ^16^. Bioluminescence imaging of the p21 promoter activity in living mice revealed that it is active at a basal level in various organs and has a tendency to increased activity in the compartments where the cells exit proliferative state and differentiate ^17,18^. Importantly, p21 plays a role in the maintenance of stem cell quiescence ^19^.

In this study, we generated and characterized a knock-in mouse model in which telomerase reverse transcriptase (TERT) is expressed from the p21^Cdkn1a^ promoter (further on p21^+/Tert^ mouse model). Since dysfunctional telomeres signal cell cycle arrest via the ATM-p53-p21 pathway ^20^, the p21^+/Tert^ model is expected to upregulate *Tert* expression in response to telomere dysfunction, thereby improving telomere maintenance. Using this unique fine-tuned regulatory loop allowing expression of telomerase specifically in cells with activated p21^Cdkn1a^ we aimed at preventing their senescence. We uncovered that p21 promoter-dependent expression of TERT decreases the number of senescent cells in the lung of old mice, promotes capillary density, and protects mice from age-associated emphysema. Single-cell transcriptomics reveal that p21-dependent expression of Tert sustains subpopulations of endothelial cells and Type II (AT2) pneumocytes providing unique insights into the mechanism by which telomerase protects against age-related emphysema.

## Results

### Generation and validation of the p21^+/Tert^ mouse model

To generate a mouse model that expresses TERT under control of the p21^Cdkn1a^ promoter, we substituted the start codon of the *Cdkn1a* gene by a *mCherry-2A-Tert* cassette (Fig. 1a). The detailed construction of the targeting vector and integration of the cassette are shown in Fig. S1a-e (see Methods). The *mCherry-2A-Tert* allele retains regulatory 5’ and 3’UTRs of the endogenous *Cdkn1a* gene. Translation of the polycistronic *mCherry-2A-Tert* mRNA produces two separate mCherry-2A and mTert polypeptides due to the ribosome skipping at the 2A sequence (Fig. 1a, Fig. S1f). The generated p21^+/Tert^ mouse therefore produces p21 protein from one allele and mCherry and Tert from the other allele. To demonstrate that TERT expressed from the *mCherry-2A-Tert* cassette is functional, we transfected Tert^-/-^ ES cells with a plasmid carrying *mCherry-2A-Tert* under control of the constitutively active pCAG promoter and checked telomerase activity *in vitro.* Telomerase activity was readily detected in multiple transfected clones (Fig. 1b). To further validate the conditional expression of the *mCherry-2A-Tert* cassette, we either irradiated p21^+/+^ and p21^+/Tert^ mice or treated p21^+/+^ and p21^+/Tert^ littermates with intraperitoneal injection of doxorubicin, known to promote p21 expression especially ^17^. We observed an increase of emitted fluorescence after whole-body ionising radiation in the p21^+/Tert^ mice after 24 and 48 hours (Fig. 1c). Along the same line, mCherry fluorescence was increased after doxorubicin treatment in the liver and kidneys from p21^+/Tert^ mice that was associated with increased levels of *mCherry-2A-mTert* transcripts (Fig. S2a, b, c). We also evaluated the level of p21 protein by semi-quantitative immunoblotting (Fig. S2C). p21^+/Tert^ mice expressed about half of the amount of p21 protein compared to p21^+/+^ in liver and kidneys pointing to the need of using use p21^+/-^ controls in subsequent experiments. Collectively, these results demonstrate that the *mCherry-2A-Tert* cassette produces active telomerase and is expressed under conditions known to induce the p53-dependent expression of p21.

**Figure 1.**
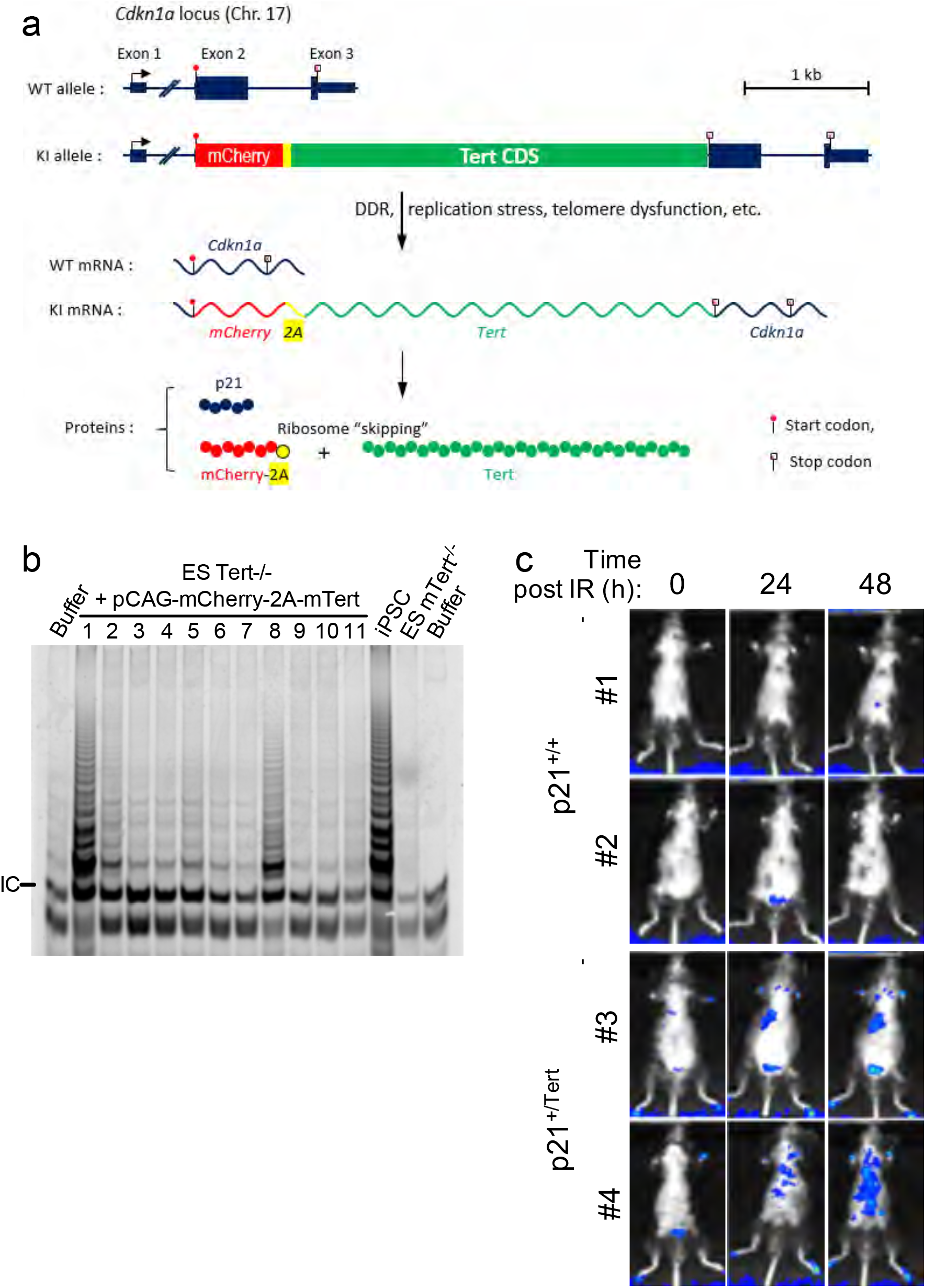
Construction and validation of the p21^+/Tert^ mouse model. **(a)** Schematic of the modified *Cdkn1a* locus, the mRNAs transcribed, and the proteins translated. The *mCherry-2A-Tert* cassette was inserted in place the start codon of Cdkn1a (see Figure S1 for details). The gene locus is drawn to scale with intron 1 contracted. The 2A peptide sequence causes a “ribosomal skip” that generates two independent polypeptides, mCherry and TERT, from the knockin (KI) allele. (**b**) Tert^-/-^ ES cells re-gain telomerase activity after transfection with the plasmid carrying mCherry-2A-Tert cassette (See Methods). *In vitro* telomerase activity was assayed using by the Telomere Repeat Amplification Protocol (TRAP). The 6 bp ladder reflects telomerase activity. The arrow indicates the internal standard control (IC). IPSC and ES Tert^/-^ cells were used as positive and negative controls, respectively. (**c**) mCherry imaging *in vivo* after whole-body ionising radiation of 1.5 gray. The fluorescence emitted by the mCherry was followed post-irradiation at the indicated times by *in vivo* mCherry imaging (Excitation = 545 nm, Background = 495 nm, Emission = 615 nm). The colour scale is indicated.

### p21-driven expression of telomerase in lungs reduces the age-related accumulation of senescent cells

To determine whether conditional expression of TERT could supress *in vivo* age-related cellular senescence in lungs, we monitored in young (4-month-old) and old (18-month-old) mice (from the 3 genotypes p21^+/+^, p21^+/-^, and p21^+/Tert^) either the number of lung parenchymal cells stained with senescence associated β-galactosidase activity (SA-β-Gal), or the number of p16-positive vascular and alveolar cells that were distinguish based on histological morphology. While young mice exhibited a very low percentage of SA-β-Gal- and p16-positive cells, lung cells from p21^+/+^ and p21^+/-^ old mice displayed a significant increase in the percentage of SA-β-Gal and p16-positive cells. In contrast, neither the percentage of SA-β-Gal, nor the one of p16-positive cells increased with age in the p21^+/Tert^ mice (Fig. 2a, b). Of note, the number of p16-marked cells was higher in vascular cells than in alveolar cells. Consistent with these results, p16 and SA-β-Gal positive lung cells were positively correlated in p21^+/+^ and p21^+/-^ cells but not in p21^+/Tert^ lung cells (Fig. 2c). We further counted the number of p21-positive cells both in alveolar and in vascular cells (Fig. 2d). The prevalence of p21 expression in vascular cells was higher than in alveolar cells. We detected a mild increase in the number p21-positive cells in old mice when compared to young mice, (Fig. 2d). In old mice, the number of p21-positive vascular or alveolar cells was not significantly reduced in the p21^+/Tert^ mice. Overall, we conclude that the p21 promoter-driven expression of TERT reduced the global number of cells either expressing p16 or positive for SA-β-gal, two robust markers of senescent cells.

**Figure 2.**
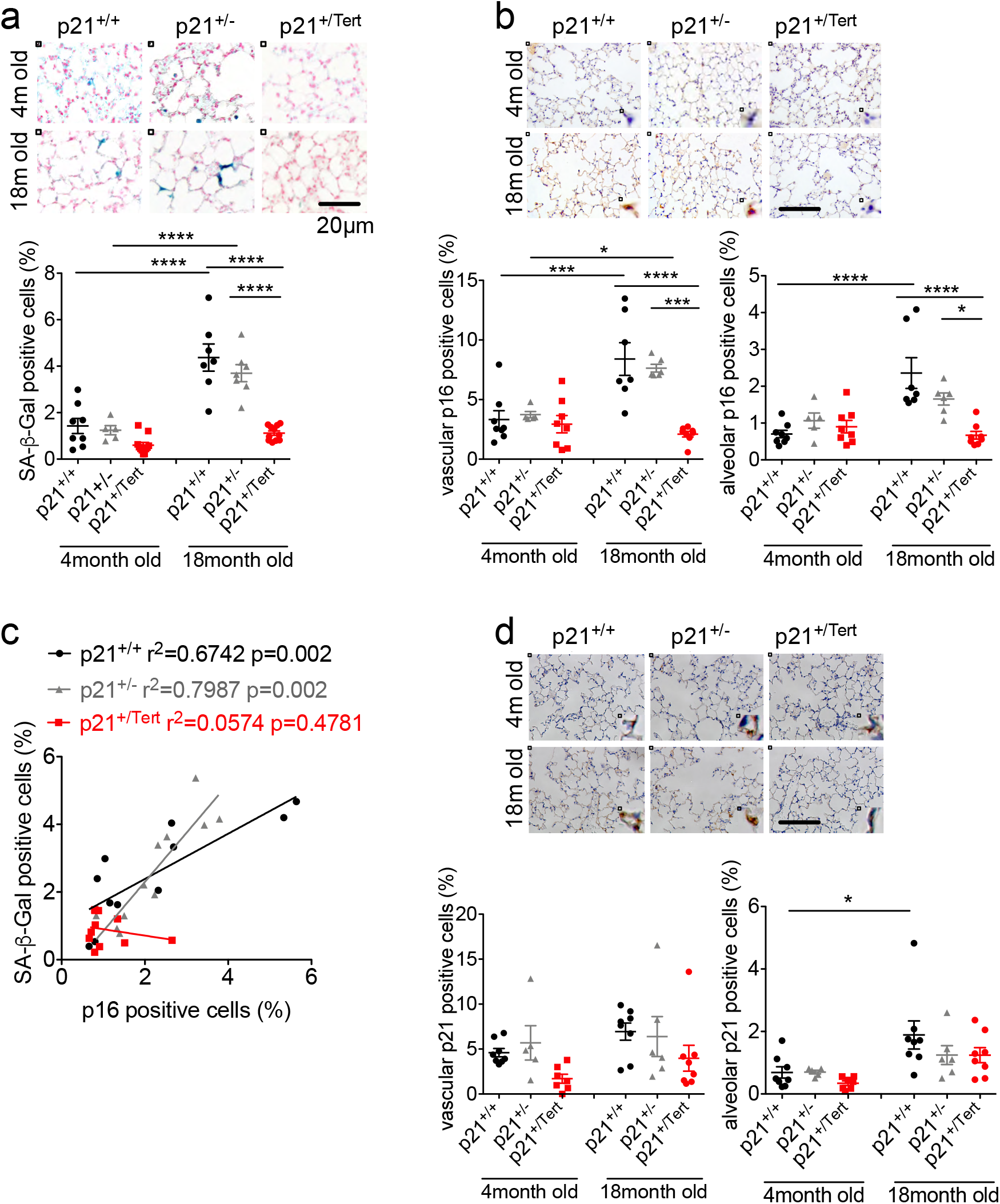
p21^+/Tert^ mice are protected against age-related associated senescent cells accumulation in the lung. (**a**) Upper panel: representative micrographs of SA-β-Gal staining in the lungs from p21^+/+^, p21^+/−^ and p21^+/Tert^ young (4-month-old) and old (18 month) mice. Blue: positive cells, Red: nuclear counterstaining using neutral red. Lower panel: quantification of the percentage of cells positive for SA-β-Gal in each group. (**b**) Upper panel: representative micrographs of mice lung immunohistochemistry against p16 (brown) with nuclear hemalum coloration (blue). Brown nuclei were considered positive. Lower left panels: Quantification of the percentage of vascular p16-positive cells. Vessels were identified according to their morphology. Lower right panel: p16-positive cells in lung tissue excluding vessels and bronchus. (**c**) Correlation between the overall percentage of p16 and SA-β-Gal positive cells in lung from mice of the indicated genotypes. Each dot represents one individual mice. (**d**) Upper panel: representative micrographs of mice lung immunohistochemistry against p21 (brown) with nuclear hemalum coloration (blue). Brown nucleus were considered positives. Lower left panel: Quantification of the percentage of vascular p21-positive cells. Lower right panel: p21-positive cells in lung tissue excluding vessels and bronchus. Scale bar is 20μm. Data are individual values +mean and SEM. Statistics: *p<0,05 **p<0,005 ***p<0,001 ****p<0,0001 using ANOVA followed by Bonferroni correction for multiple tests. Not significant differences are not displayed.

We thus investigated the presence of very short telomeres (VST) in lungs from six p21^+/+^ and six p21^+/Tert^ old mice compared to young counterparts by using the TeSLA method that allows measuring the distribution of the shortest telomeres in cells and tissues ^21^ (Fig. S3a, b, c). Quantification of the results (See Methods) indicated that the fraction of short telomeres was not significantly different in between p21^+/+^ and p21^+/Tert^ mice (Fig. S3a). However, the cumulative number of VST telomeres in lung cells from p21^+/+^ mice was highly correlated with the number of cells endowed with SA-β-Gal activity (Fig. S3b). This correlation was also observed at a lower level in p21^+/Tert^ mice suggesting that the fraction of very short telomeres is a marker of senescence in mice (that are otherwise TERT^+/+^).

### Smooth muscle cells from p21^+/Tert^ pulmonary arteries bypass senescence ex-vivo

To deeply characterize the p21-promoter dependent expression of TERT on cellular senescence, we sought to test ex-vivo its effect on the proliferation of cultured pulmonary artery smooth muscle cells (PA-SMCs) isolated from lungs of young mice. PA-SMCs of the three genotypes (p21^+/+^, p21^+/Tert^ (littermate) and p21^+/−^) were cultured in standard oxidative stress conditions (20% oxygen) for which we expect TERT to be induced in most of the cells (Fig. 3a, b). Cells from p21^+/+^ and p21^+/-^ mice entered into senescence after a few passages, with a final mean population doubling level (PDL) of 15,51 (+/− 4,14) and 36,09 (+/− 8,30) respectively. p21^+/Tert^ PA-SMCs on their side mice never entered into senescence and proliferated with a much faster rate (Fig. 3a). The difference in cumulative PDL between p21^+/Tert^ and both p21^+/+^ and p21^+/−^ became significant at passage 7. We confirmed the expression of the mCherry-2A-Tert cassette by qRT-PCR (Fig. 3c), and also detected a slight increase of telomerase activity at passages 2-4, before PA-SMCs cumulative PDL curve became significantly different (Fig. 3d). We next measured the load of VSTs in p21^+/+^ and p21^+/Tert^ PA-SMCs (Fig. 3e). While the cumulative number of short telomeres increased with passages in p21^+/+^ PA-SMCs, the load of short telomeres was identical at passages 4 and 10 in p21^+/Tert^ PA-SMCs (Fig. 3e). Therefore, proliferation of the cultured p21^+/Tert^ PA-SMCs coincides with conditional expression of TERT and reduced prevalence of the VSTs suggesting that very short telomeres could be repaired by telomerase in mice ^22^.

**Figure 3.**
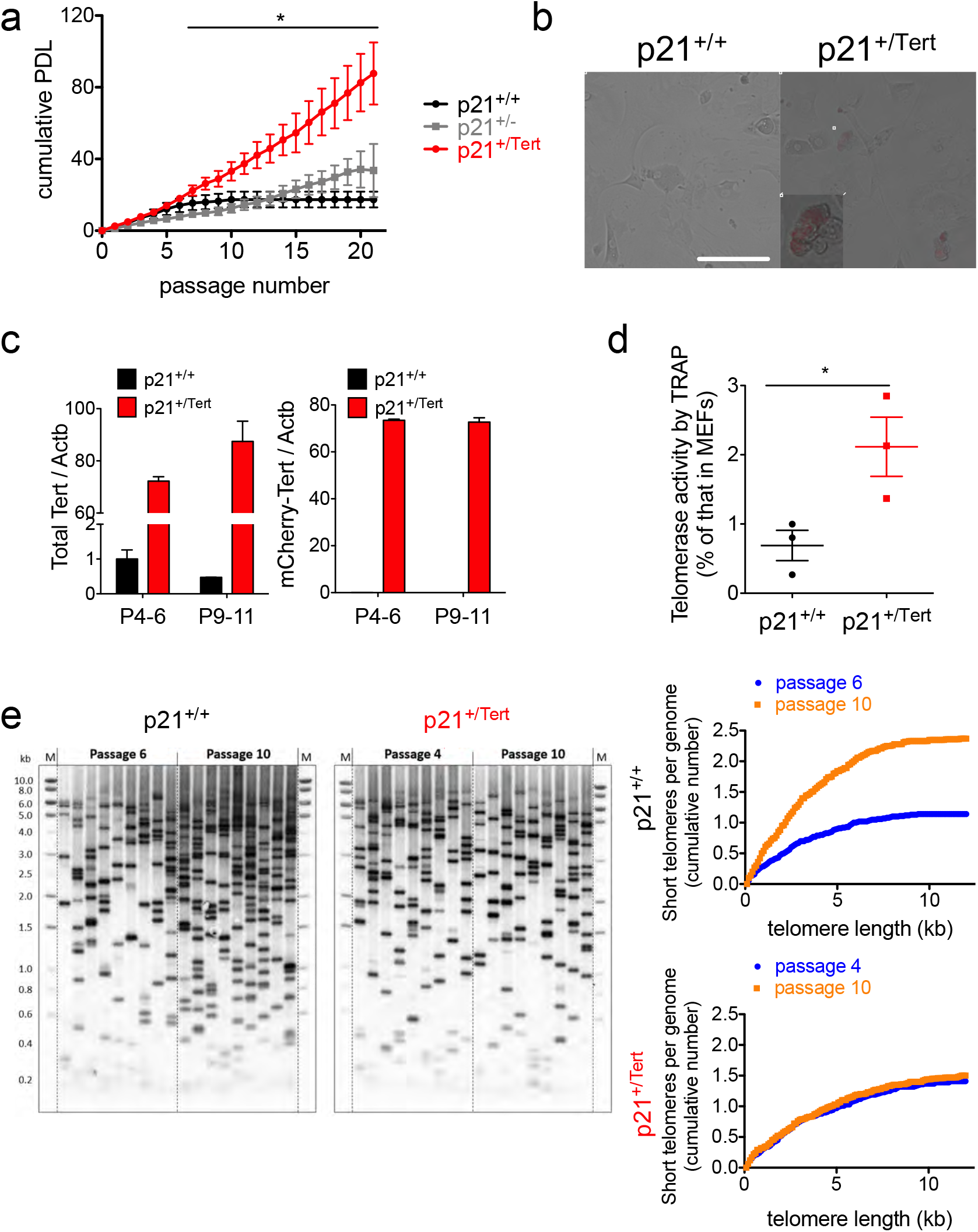
p21-promoter dependent TERT expression bypasses senescence in PA-SMCs ex-vivo. **(a)** Cumulative population doubling level (PDL) of pulmonary-artery smooth muscle cells (PA-SMCs) isolated from mice from the three mentioned genotypes. The data points are the means of 8 independent cultures established from individual mice. *p<0,05 comparing p21^+/Tert^ with p21^+/+^ and p21^+/−^, student t-test. **(b)** PA-SMC from p21^+/+^ (left) and p21^+/Tert^ (right) littermate mice were observed under the microscope at passage 4. Representative fields were obtained using transmitted light and mCherry filter. Higher magnification of a mCherry-positive cell is shown in the bottom left. Scale bar is 100 μm. (**c**) Quantification of the *Tert* mRNA levels in PA-SMCs from the p21^+/+^ and p21^+/Tert^ 4-month-old mice (littermates) at early (p4-6) and late (p9-11) passage. Left panel represent the level of endogenous Tert mRNA, right panel represents the level of KI allele. Nearly all *Tert* mRNA is transcribed from the KI allele. The means of three independent measurements are plotted, and the error bars are SEs. (**d**) Telomerase activity measured by qTRAP at early passages. The data points correspond to vascular PA-SMC cultures established from individual p21^+/+^ and p21^+/Tert^ 4-month-old mice. * p <0.05 from the two-sided *t* test. (**e**) Analysis of the short telomere fraction by Telomere Shortest Length Assay (TeSLA) in the cultured PA-SMCs from p21^+/+^ and p21^+/Tert^ mice. Left panels depict Southern blots probed for the TTAGGG repeats, and the right panels show quantification of the cumulative number of short telomeres across the telomere length thresholds.

### Conditional expression of telomerase protects against age-related emphysema

We next sought to determine the impact of the conditional expression of telomerase on age-related manifestations in the lung, such as emphysema as it has been widely documented in old C57Bl6 mice ^23^. Emphysema is characterized by lung airspace enlargement as a consequence of a decrease of lung elasticity with advanced age ^24^. Morphometry studies ^25^ did not reveal air space enlargement in young mice from the three genotypes (Fig. 4a). Old p21^+/+^ and p21^+/−^ mice developed emphysema reflected by an increase of alveolar size (measured by the mean-linear intercept or MLI). In contrast, old p21^+/Tert^ mice did not exhibit air space enlargement (Fig. 4a).

**Figure 4.**
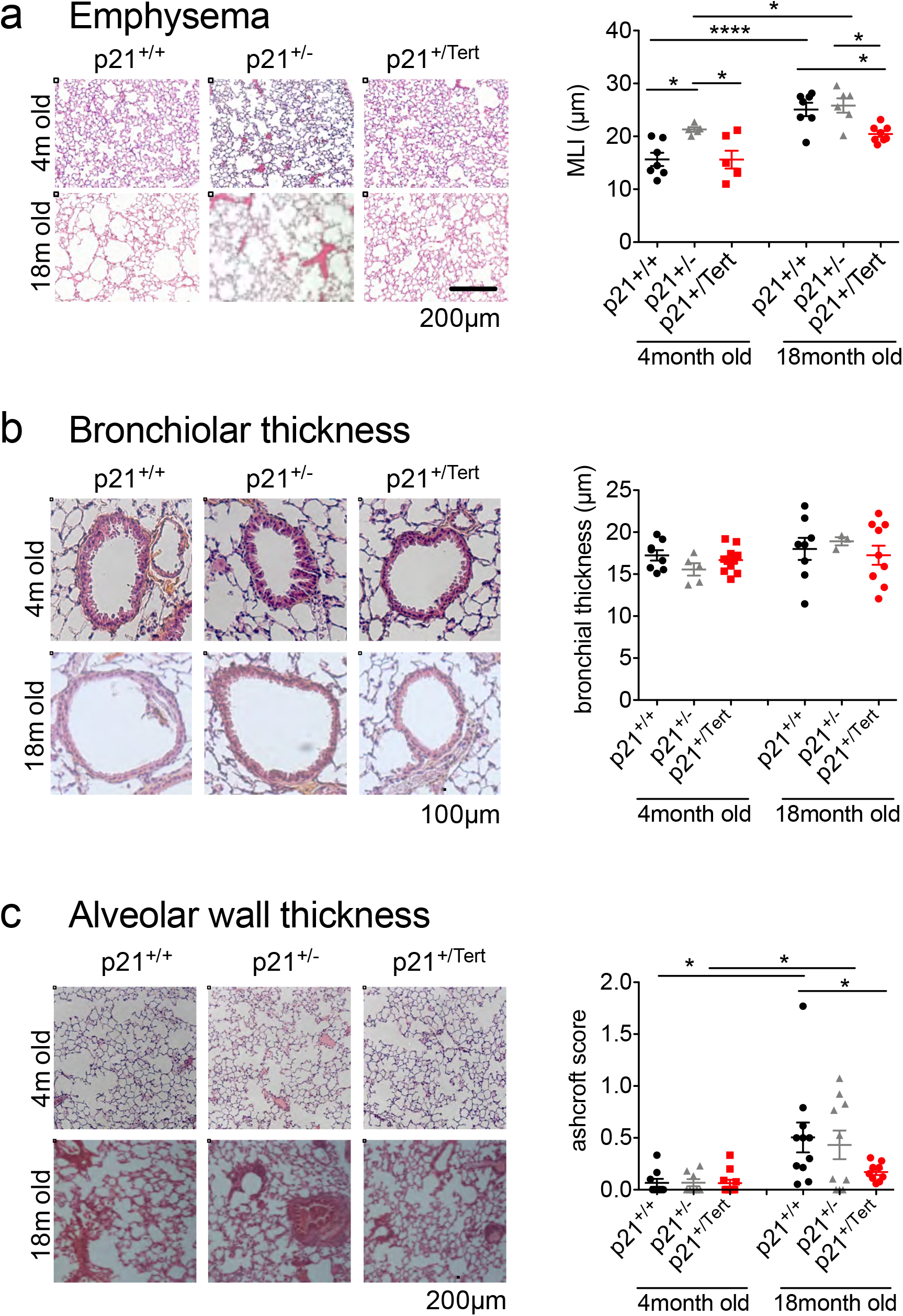
Tert expression under the control of the p21 promoter protects against age-related lung emphysema and fibrosis. (**a**) Emphysema assessment is obtained by measuring the mean linear intercept (MLI) in lungs from 4 and 18-month-old p21^+/+^, p21^+/Tert^ (littermates), and p21^+/−^ mice. Scale bar is 200μm. (b) Bronchiolar thickness assessment is obtained by measuring the epithelial wall thickness in lungs from young and old mice from the three genotypes. Scale bar is 100μm. (c) Lung fibrosis assessment is obtained by histopathological analysis using modified Ashcroft scoring on young and old mice from the three genotypes. Scale bar is 200μm. (a, b, c): Data are individual values +mean and SEM. Statistics: *p<0,05 **p<0,005 ***p<0,001 ****p<0,0001 using ANOVA followed by Bonferroni correction for multiple tests. Not significant differences are not displayed.

We also measured bronchiolar thickness and sought for manifestations related to lung fibrosis in the same cohort of mice. It is known that collagen deposition in the lung increases with age ^26^. These analyses did not reveal significant variation of bronchiolar wall thickness with age or between genotypes (Fig. 4b). In contrast, we observed a discreet but significant increase of the Ashcroft score (a morphological test for pulmonary fibrosis) in lungs from p21^+/+^ and p21^+/−^ mice with age, but not in p21^+/Tert^ mice (Fig. 4c), indicating a minimal alveolar fibrous thickening in p21^+/+^ and p21^+/−^ mice with age. Taken together, these results indicate that p21-dependent expression of TERT protects mice from age-related emphysema and morphological changes of the lung.

### Suppression of emphysema in p21^+/Tert^ lungs is associated with endothelial cell proliferation and capillary density

We next sought to determine whether this lung protection was associated to higher levels of cellular proliferation, which could then favour cell renewal and preserve tissue architecture with age. We assessed cell proliferation in the lungs of old mice from the three genotypes by analyzing BrdU incorporation (see Methods). We found a higher cell proliferation level in p21^+/Tert^ mice compared to control mice (Fig. 5a) that was associated to lower MLI (Fig. 5b, c) and reduced number of senescent cells (Fig. 5d). Since BrdU-positive cells were frequently found in the vascular compartment, we wondered if this enhanced proliferation capacity was associated with a preserved vasculature in p21^+/Tert^ mice. We thus measured capillary density by anti-isolectin B4 immunofluorescence ^27^. We observed a strong tendency for increased capillary density with age in the p21^+/Tert^ (Fig. 5e). These results suggest that TERT promotes endothelial cell proliferation with age thus preserving capillary density.

**Figure 5.**
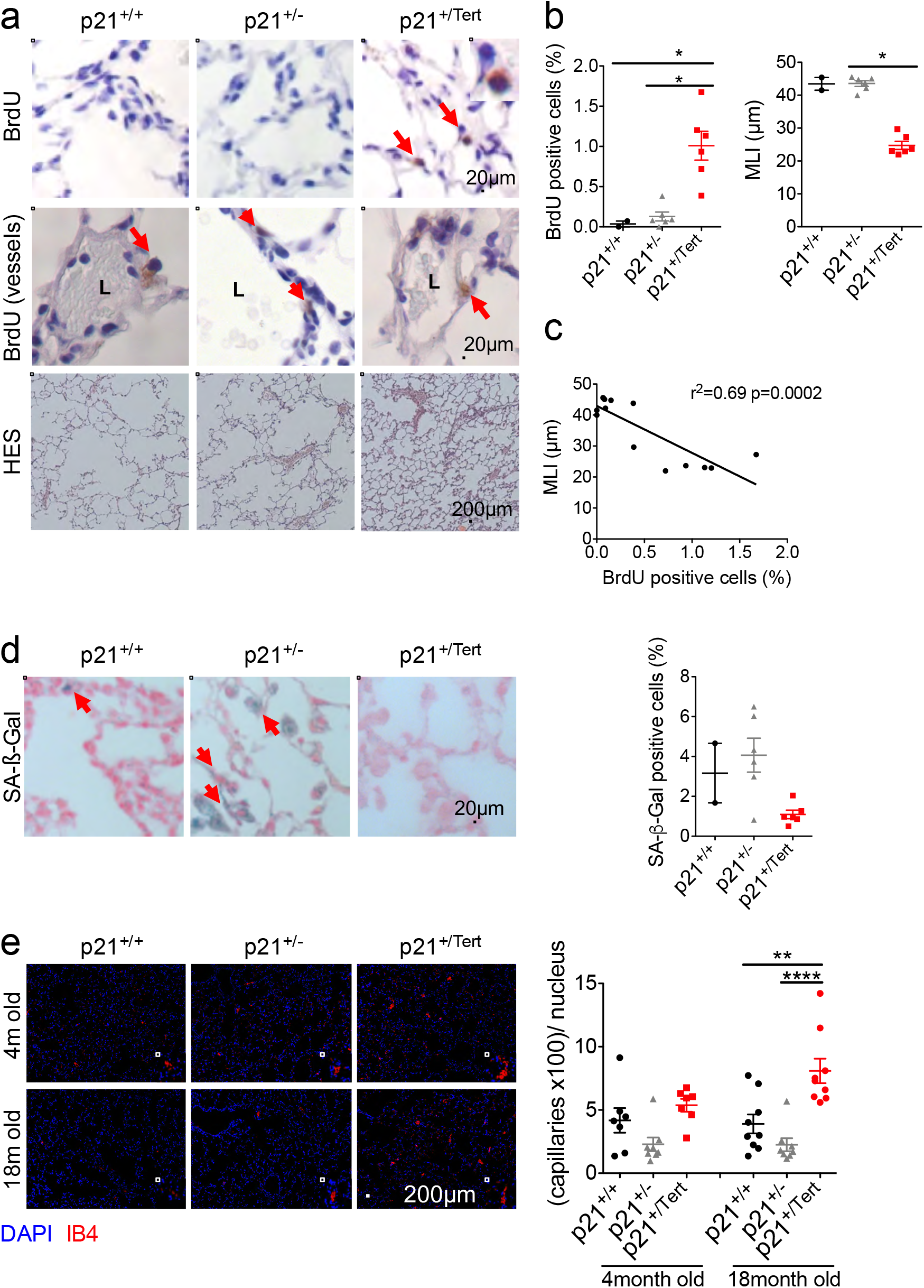
Suppression of emphysema in p21^+/Tert^ lungs is associated with vascular cell preservation. 18-month-old mice were injected with Bromodeoxyuridin (BrdU) intra-peritoneally 24h before organ sampling. (**a**) BrdU immunostaining was performed using anti-BrdU antibody in lungs from p21^+/+^ and p21^+/Tert^ littermates mice along with p21^+/−^ mice (upper panel). Positive cells are stained in brown (red arrows). Nucleus were stained using hematoxillin (blue). Middle panel emphases on vessels where most of the BrdU-positive cells were found. L= Vessel lumen. Lower panels show haematoxylin-eosin-safran (HES) staining of the lungs for emphysema assessment. (**b**) Quantification of BrdU-positive cells in lungs expressed in percentage for the overall lung tissue (left panel) and emphysema assessment by measurement of the mean linear intercept (MLI, in μm). (**c**) Correlation of the percent of BrdU positive cells and emphysema for each mouse. (**d**) Measurement of the percentage of SA-β-Gal positive cells (in blue). Left panel: Representative staining of positive cells (red arrow). Right panel: percentage of SA-ß-Gal positive cells for each mouse. (**e**) Capillary density in lung was assessed in p21^+/+^, p21^+/−^ and p21^+/Tert^ mice aged of 4 and 18-months. Immunofluorescence experiment was performed by using anti-isolectinB4 antibody (red) and DAPI staining (blue). Higher magnification examples of capillaries are presented on the lower right corner of each field. Scale bar is 200μm. Right panel: number of capillaries normalized by the number of DAPI-positive cells. Data are individual values +mean and SEM. Statistics: *p<0,05 **p<0,005 ***p<0,001 ****p<0,0001 using ANOVA followed by Bonferroni correction for multiple tests. Not significant differences are not displayed.

### Protection against age-related emphysema requires the catalytic activity of TERT

To determine whether the catalytic activity of telomerase was required for lung protection, we created a new mouse line (called p21^+/Tert-CI^) based on p21^+/Tert^ mice, but carrying a point mutation (D702A) in the active site of TERT ^28^. We performed the experiments described above (MLI, cellular senescence, cell proliferation and capillary density) in p21^+/Tert^ and p21^+/Tert-CI^ mice at the age of 18 month. While values for p21^+/Tert^ were comparable to those obtain in previous experiments, we found that p21^+/Tert-CI^ presented a significant increase in MLI (Fig. 6a), associated with an increase in both the number of vascular and alveolar p16-positive cells (Fig. 6b) and SA-β-Gal staining (Fig. 6c). p21^+/Tert-CI^ mice also presented an increase in vascular p21-positive cells compared to p21^+/Tert^, albeit not in alveoli (Fig. 6d). We also observed a very low level of proliferative cells in p21^+/Tert-CI^ mice lungs compared to p21^+/Tert^ mice (Fig. 6e). Finally, capillary density in p21^+/Tert-CI^ mice was also significantly decreased with respect to p21^+/Tert^ mice (Fig. 6f). Overall, p21^+/Tert-CI^ mice exhibit the same aging characteristics in the lung as control mice. These results strongly support the idea that TERT acts either by elongating critically short or damaged telomeres ^22,29,30^.

**Figure 6.**
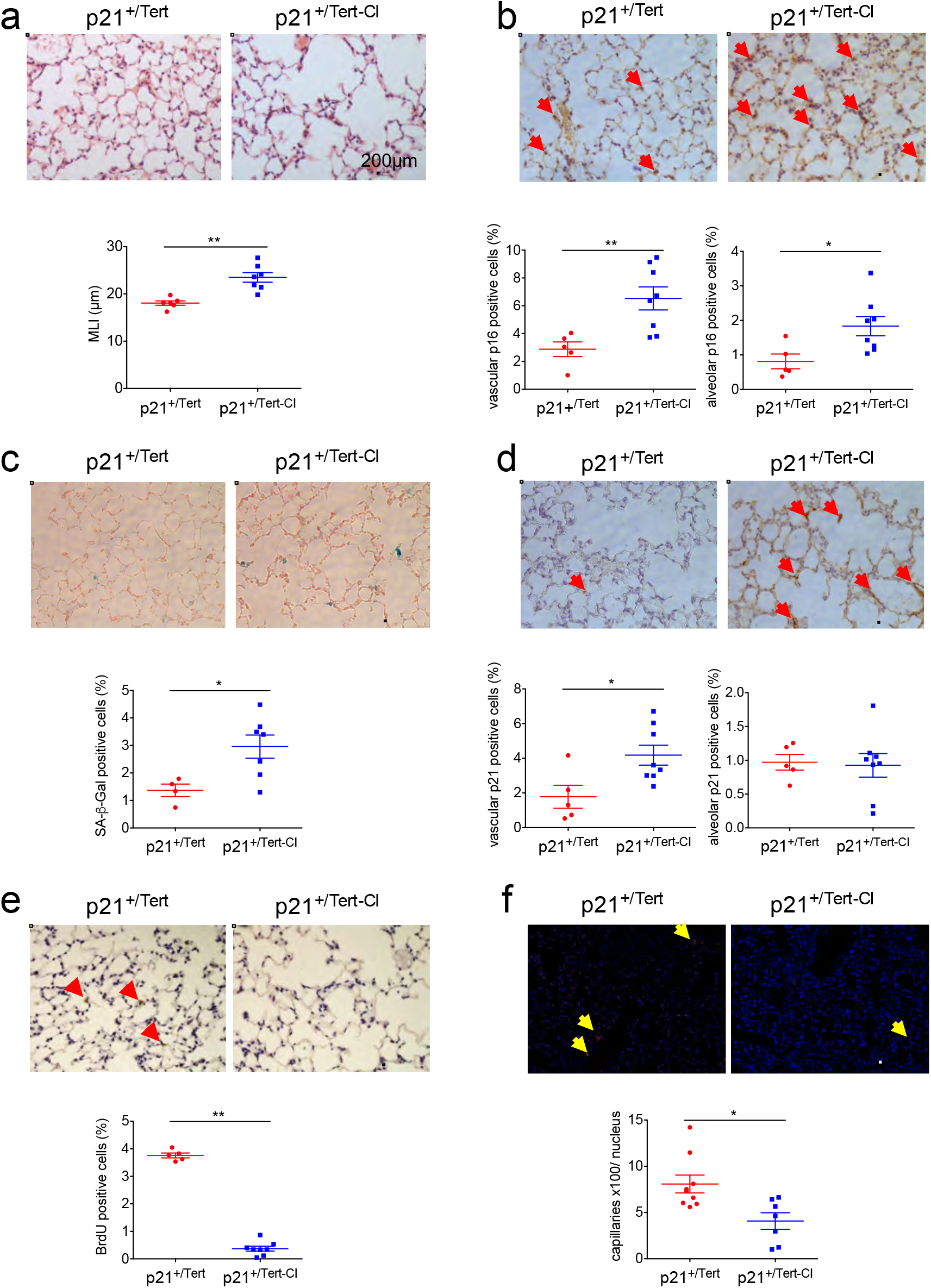
Emphysema, cellular senescence, and capillary density in old mice expressing a catalytically inactivated TERT (p21^+/Tert-CI^ mice) 18-month-old p21+/Tert and p21+/Tert-CI mice were injected with Bromodeoxyuridin (BrdU) intraperitoneally 24h before organ sampling and sacrificed. Upper panels are representative micrographs, lower panels are quantifications. (**a**) Emphysema was assessed by measuring the mean linear intercept (MLI, in μm) on haematoxylin-eosin-Safran stained lungs. (**b, d, e**) Immonuhistochemistry was performed using anti-p16 (B), anti-p21 (D), and anti-BrdU (E) antibodies (red arrows) with nuclear hemalum counter-staining (blue). Results are expressed as percentage of vascular cells (left) or percentage of alveolar cells (right). (**c**) SA-β-Gal staining (blue) with nuclear neutral red counter-staining (red). Results are presented as overall percent positive cells. (**f**) Capillary density was assessed using anti-isolectinB4 antibody (red) with DAPI (blue). Yellow arrows indicate positive signal. Results are presented as the number of capillaries normalized by the number of DAPI-positive cells. Data are individual values +mean and SEM. Statistics: *p<0,05 **p<0,005 using Mann-Whitney test. Not significant differences are not displayed. Scale bar is 200μm.

### A subcluster of p21^+/Tert^ gCap cells express markers associated with regeneration

To further understand how ectopic expression of telomerase driven by the p21 promoter could suppress emphysema in old mice, we performed single-cell transcriptomics (using 10x Genomics Chromium system) on lung cells from 18 month-old p21^+/+^, p21^+/Tert^ and p21^+/−^ mice. For a single lung, around 5000 individual lung cells were encapsulated and libraries were prepared and sequenced. We performed dimensionality reduction and unsupervised cell clustering to identify distinct cell types based on shared and unique patterns of gene expression (see Methods). For each mouse, clustering of gene expression matrices identified cell types that were in good agreement with the published Mouse Cell Atlas (MCA) ^31^ (Fig. S4a, Table 1). Cell-type classes of immune and non-immune cells were similar for the 3 mice (Fig. S4a, b). We found that Cdkn1a (p21) was mainly expressed in interstitial and alveolar macrophages, monocytes, and in dendritic cells while in resident lung cells, it was mainly expressed in endothelial cells (Fig. S4c).

Because pulmonary endothelial cells (EC) seem to play a key role in the development of emphysema, we first targeted our single-cell transcriptomics analysis to EC (Fig. S5C). We annotated lung EC to specific vascular compartment based on recently published markers of lung EC ^32^ (Fig. S5). We identified 5 classes of EC shown in Fig. 7a,b (Table 2). Interestingly, capillary class 1 and 2 were recently identified as aerocytes (aCap) involved in gas exchange and trafficking of leucocytes and as general capillary cells (gCap) that function in vasomotor tone regulation and in capillary homeostasis ^33^. We reasoned that p21-expression should reflect Tert expression in p21^+/Tert^ cells since both genes belong to the same polycistronic mRNA. For each mouse (p21^+/+^, p21^+/−^, p21^+/Tert^), we compared differentially expressed genes (DEG) between p21-positive and -negative EC. We uncover that a number of genes were significantly overexpressed specifically in p21-positive cells in p21^+/Tert^ EC (Fig. 7c, Fig. S5) Strikingly, most of these genes are functionally related (Fig. 7d) and are positively regulated by the stress response gene Atf3 (Cyclic AMP-dependent transcription factor 3) involved in EC regeneration ^34^ (See discussion).

**Figure 7.**
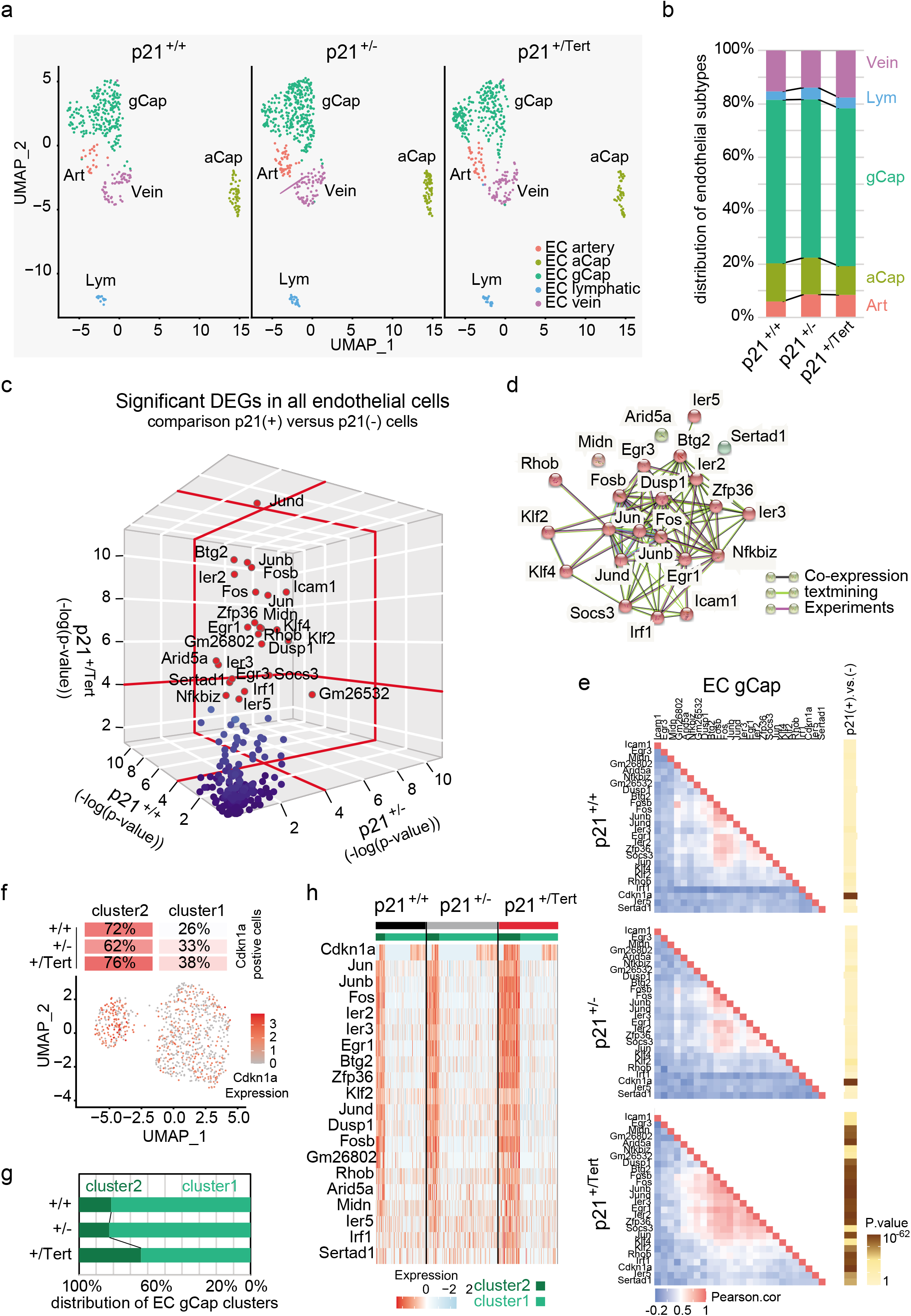
A subclass of p21^+/Tert^ endothelial capillary cells shows a transcriptomic signature associated with EC regeneration. (**a**) UMAP clustering of lung endothelial cells from WT, p21^+/−^ and p21^+/Tert^ mice. Cell populations were identified based known markers for these endothelial cell subtypes^32^ (Fig. S5). (Art) artery EC cells; (Cap_1; also called aCap^33^) capillary 1 EC cells; (Cap_2; also called gCap^33^) capillary 2 EC cells ; (Lym) lymphatic EC cells and (Vein) vein EC cells. (**b**) Distribution of EC subtypes in p21^+/+^, p21^+/−^ and p21^+/Tert^ mice in percentage. (**c**) 3D plots of the significant p-values (-log10 p > 1.42) for the DEGs between p21-positive (Cdkn1a > 0) and p21-negative (Cdkn1a = 0) EC from lungs from p21^+/+^, p21^+/−^, and p21^+/Tert^. Genotypes are indicated on the X, Y, and Z axis. The highly significant genes (-log10 p > 4) are shown as red and green dots. (**d**) Predicted interactions genes identified in (C). The interaction network was downloaded from the STRING database. (**e**) Triangular heatmap representing the pairwise correlation coefficients (pearson correlation) for the DEG in gCap. Genotypes are indicated in the figure. Genes are ordered according to a hierarchical clustering. Correlation coefficient values are represented according to the indicted colour scale. Coloured rectangles on the right indicate the p.value for each gene after comparison of their expression in p21 positive (Cdkn1a > 0) versus negative (Cdkn1a = 0) gCap cells. (**f**) UMAP clustering of gCap cells (merge of the 3 genotypes). Note the clear demarcation of two sub clusters, characterized by the number of cells expressing Cdkn1a (p21). (**g**) % of gCap cells belonging to cluster 1 and 2 for the indicated genotypes (**h**) Heatmap of DEG in in gCap cells. The dark and light green bars represent cells from cluster 2 and 1, respectively.

Because, old p21^+/Tert^ mice had an increased lung capillary density, we focussed our attention to capillary class 2 cells (gCap) ^33^. We uncover that many genes overexpressed in p21-positive (versus – negative) EC were also significantly expressed in p21^+/Tert^ gCaps (Fig. S6). For the 3 mice, we performed Pearson correlations to analyse co-expression in gCaps of the genes up-regulated in p21-positive cells (Fig. 7e). Remarkably, we found that the functionally related genes shown in Fig. 7D were significantly co-expressed in p21^+/Tert^ gCaps suggesting that these genes could be directly or indirectly co-regulated by Tert. We further uncover that gCaps could be subclustered in two clusters named 1 and 2 (Fig. 7f). Of note, cluster 2 was overrepresented in p21^+/Tert^ gCaps raising the possibility that the differences in gene expression could be related to the increased number of cells belonging to cluster 2 (Fig. 7G). Moreover, we found that most FRG that we defined above were mainly overexpressed in p21-positive cluster 2 cells in comparison to cluster 1 cells (Fig. 7h).

Collectively, these results suggest that TERT sustain the growth of a subpopulation of gCap cells (cluster 2) expressing markers associated with EC regeneration (see discussion).

### A subclass of p21^+/Tert^ AT2 cells express markers of club cells and S/G2M associated genes

Because Tert targeting in AT2 cells was shown to prevent pulmonary fibrosis progression ^35^, we next clustered p21^+/+^, p21^+/−^, and p21^+/Tert^ AT2 cells resulting in a detailed map of AT2 cells for each genotype. Two-dimensional representations revealed 3 main clusters within AT2 cells whose proportion differed for the three mice (Fig. 8a). Cluster 1 was found in all three mice, with an overrepresentation in WT mice. Cluster 2 was strictly specific for p21^+/−^ and p21^+/Tert^ mice while cluster 3 was over-represented in p21^+/Tert^ mice.

**Figure 8.**
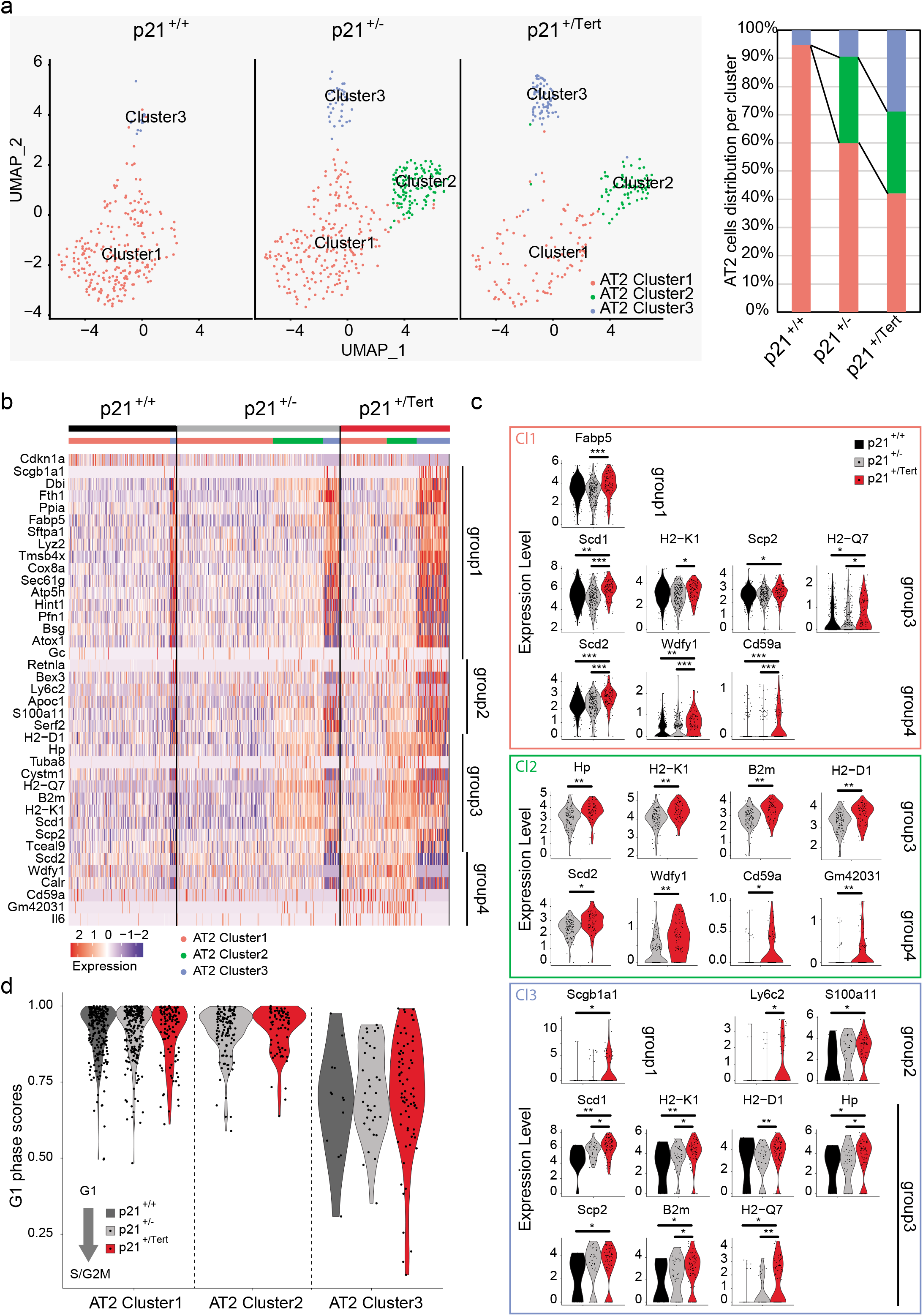
p21^+/Tert^ AT2 cells are enriched in a subclass of AT2 cells expressing markers of epithelial cells sustaining alveolar regeneration. **(a)** UMAP showing subclusters of AT2 cells. Right panel represents AT2 cell distribution within each cluster. **(b)** Heatmap of differentially up-regulated expressed genes (upDEGs) in p21^+/Tert^ AT2 single cells compared to WT and p21^+/−^. Columns represent individual cells grouped by cluster. Light red, green, and blue bars delineate cells from cluster 1, 2 and 3, respectively. Genes are grouped regarding to their expression levels in each cluster (hierarchical clustering on the Pearson correlation between genes). The colour code represents the z-score of expression. (**c**) Violin plots show gene expression (log2 scale) of only significant upDEGs in single-cell p21^+/Tert^ (red) compare to WT (black) or p21^+/−^ (grey) within each cluster. Dots represent individual cells. Wilcoxon rank-sum test with continuity correction was performed to test significance (*P < 0.05; **P < 0.01; ***P < 0.001). (**d**) Violin plots show G1-phase score per cell based on the expression of cell cycle-specific markers. AT2 cells are spited according to cluster and genotype (WT (black); p21^+/^ (grey) or p21^+/Tert^ (red)).

We analysed DEGs in between AT2 cells from p21^+/+^, p21^+/−^, and p21^+/Tert^ mice. A large number of genes were differentially expressed in p21^+/Tert^ AT2 cells compared to either p21^+/+^ or p21^+/−^ (Table 3). However, a number of these genes were also differentially expressed in between p21^+/−^ and p21^+/+^ likely reflecting differences in cell clusters in between the three mice. Hierarchical clustering of genes allowed the identification of 4 groups of genes that were up-regulated in p21^+/Tert^ AT2 cells (Fig. 8b, Fig. S7). The first group of genes was mainly overexpressed in AT2 cluster 3 whose number of cells was increased in p21^+/Tert^. This group was in particular characterized by the expression of the club cell markers Scgb1a1 (Secretoglobin family 1A member 1) as well by the virtual lack of Cdkn1a expression (Fig. 8b). Genes from group 2 and 3 were mainly overexpressed in AT2 cluster 2 and 3 (Fig. 8b). We could identify in the group 3 a signature of functionally related genes containing MHC class 1 genes (H2-D1, H2-K1, H2-Q7, B2m) and Scd1 (Acyl-CoA desaturase) (see discussion). In contrast to the other groups, genes from group 4 were not expressed in AT2 cluster 3. Individual expression of representative genes is shown in Fig. 8c. Interestingly, we uncovered that a higher number of cells belonging to cluster 3 cells (compared to cluster 1 and 2 cells) expressed genes associated with the S/G2M phase when compared to cluster 1 and 2, which mainly expressed G1-phase genes (Fig. 8d), suggesting that cluster 3 may support AT2 cell regeneration. Many more genes were significantly down-regulated in p21^+/Tert^ AT2 cells compared to p21^+/+^ or p21^+/−^ likely reflecting differences in the distribution of the AT2 cell classes in p21^+/Tert^ compared to p21^+/+^ and p21^+/−^ (Fig. S7).

In summary, p21^+/Tert^ AT2 cells are enriched in a subclass of AT2 cells (cluster 3) that are characterized by markers of club cells and expression of S/G2M genes while p21 haploinsufficiency promotes the generation of cluster 2 cells. Cluster 3 shares features with cells involved in the repair of alveolar epithelia after lung injury (see discussion).

## Discussion

Numerous studies have shown that abnormally short telomeres and senescent cell accumulation are found in the lungs of COPD patients suggesting a direct relationship between short telomeres, senescence and the development of the disease ^47,36–38^. These conclusions have been strengthened by mouse studies showing that alveolar stem cell failure is a driver of telomere-mediated lung disease ^5^. Collectively these results suggest that DNA damage, cellular senescence, stem cell exhaustion, and likely mitochondrial dysfunction, all induced by accumulation of short telomeres, contribute together to lung injury. Because TERT may reverse these processes, TERT-based gene therapy ^39^ may be clinically beneficial to treat COPD. However, the mechanism by which *in vivo* telomerase expression could prevent the occurrence of emphysema during aging remains to be elucidated.

In this study, we designed a fine-tuned regulatory loop allowing the conditional expression of telomerase under conditions that induce the expression of p21, including the accumulation of critically short telomeres. Importantly this conditional expression of TERT has been introduced in telomerase-positive mice (Tert^+/+^), e.g. with long telomeres. Although p21 expression is not exclusively dependent of p53, the p21^+/Tert^ mouse provides a unique tool allowing us to evaluate the impact of TERT re-expression on age-related diseases. We exploited the fact that C57BL/6NRj mice naturally develop age-related emphysema to test *in vivo* the effects of TERT specific expression.

We report that the p21 promoter-driven expression of TERT in old lungs reduces the number of cells expressing robust markers of senescence such as p16 and β-gal. This is not due to the haploinsufficiency of p21 since the reduction of senescent cells was not observed in p21^+/−^ mice, although we cannot completely exclude a role of the reduced level of p21 in the phenotypes of the p21^+/Tert^ mouse. p21^+/Tert^ mice were protected from age-related emphysema and mild fibrous alveolar thickening naturally occurring in p21^+/+^ and p21^+/−^ old mice. Because, we were able to correlate the presence of very short telomeres (<400 bp) in lung cells with cellular senescence, we assume that the p21-induced Tert might repair damaged telomeres. Indeed, it has been previously reported that the DNA damage response at telomeres was contributing to lung aging and COPD ^29^.

Remarkably, we further uncovered that old p21^+/Tert^ mice exhibited an increased proliferation of cells compared to control mice. In agreement with this observation, capillary density was increased in p21^+/Tert^ mice indicating that p21-promoter dependent expression of TERT during aging improves capillary vasculature maintenance. These results suggest that TERT could prevent age-related emphysema by promoting microvasculature. This hypothesis is supported by previous studies showing a link between vascular density, endothelial cells and emphysema ^40–42^.

The single-cell experiments provide a theoretical framework that potentially explains how telomerase expression promotes capillary density in old mice. We discover that p21-positive p21^+/Tert^ EC express a signature of functionally related genes (Fos, Junb, Fosb, Nfkbiz, Jund, Dusp1, Jun, Rhob, Egr3, Btg2 Egr1, Zfp36, Ier2, Ier3, Ier5, Klf2, Klf4, Socs3, Icam1, Irf1), many of them being positively regulated by Atf3, an important transcriptional regulator required for regeneration of the endothelial lining of large arteries following injury ^34^. We further report that a subpopulation of gCap cells was overrepresented in p21^+/Tert^ gCap cells and preferentially expressed the regeneration signature. These results suggest that stress response genes related to Atf3 that contribute to repair large vessel injury ^34^, may also promote an increase of capillary density. Among these genes, we found AP-1 factors (Fos, Fosb, Jun, JunB, JunD) that are involved in numerous cellular functions such as proliferation, survival, transformation, and protection against oxidative stress (Reddy and Mossman, 2002). FRG also include the transcription factor Egr1 that was shown to play an important role in the formation of new blood vessels from the pre-existing vasculature ^45^, and Klf2 (Krüppel-like factor 2) that regulates endothelial gene expression in response to shear stress due to flow ^46^. This signature also includes a number of genes attenuating inflammation such as Nfkbiz ^47,48^, Zfp36 ^49,50^, Socs3 and Dusp1 ^51^. We propose from these results that gCap cluster 2 cells by expressing a specific signature associated with EC regeneration could account for the increase capillary density that we observed in p21^+/Tert^ mice and therefore contribute to the suppression of emphysema in p21^+/Tert^ _mice._

We also focussed our single-cell studies to Type II alveolar cells since their dysfunction has been linked to lung disease ^52,53^. We uncover that p21^+/Tert^ AT2 cells were enriched in a specific subpopulation that expressed several markers of club cells and characterized by the lack of expression of Cdkn1a. Remarkably, these cells exhibited a higher number of dividing cells. These results therefore raise the possibility that telomerase expression may promote the mobilization of a source of club cells that differentiates into AT2 cells while retaining markers of the original lineage. This cluster (3) of AT2 cells shares features with cells involved in the repair of alveolar epithelia after lung injury ^54,55^ but are different from bronchioalveolar stem cell (BASC) ^56–58^. Strikingly, cluster 2 and 3 cells overexpressed MHC class I genes. These genes were reported to mark a minor subpopulation of club-like cells endowed with the ability to support alveolar regeneration in mice after bleomycin injury ^59^. Interestingly, the four MHC class I genes H2-K1, H2-Q7, H2-D1, and B2m (whose high expression is usually associated with an interferon-gamma signature) together with Acyl-CoA desaturase 1 (Scd1) were reported to be overexpressed in AT2 cells from old mice compared to young mice ^60^.

We found that inactivation of telomerase catalytic activity seemed to abolish the protection conferred by the p21^+/Tert^ cassette pointing out the crucial role of TERT canonical activity in sustaining lung function. These results support studies indicating that short telomeres are important for the predisposition to lung disease in patients with mutations affecting telomere elongation by telomerase ^7^. Our results support a model in which mice accumulate with age a subset of presenescent cells with critically short/damaged telomeres that are not yet engaged into an irreversible senescence process. The conditional expression of TERT by elongating/repairing the very short telomeres may prevent these cells to initiate a DNA damage response and to become senescent. This process would be particularly important in maintaining the regenerative capacities of stem/progenitor cells involved in sustaining the alveolar function as discussed above and as proposed by ex-vivo experiments by ^4^. Overall, our results shield new light for the development of TERT-AAV9 treatments to treat lung diseases ^35^.

## Acknowledgments

We are grateful to the CRCM animal facility for taking care of the mouse strain colonies, to Manon Richaud from the CRCM flow cytometry and cell-sorting platform, and to Lionel Spinelli, Arnaud Guille and Samuel Granjeaud for useful discussions and technical support for the single-cell analysis. We thank Lea Harrington for providing mTert-/- mouse ES cells.

Work in VG’s Lab is supported by “La Ligue Contre le Cancer”, Equipe Labellisée, La Région SUD (Volet Général), The Canceropole PACA (Projet Emergent), the “Institut National du Cancer” (INCA), PLBIO 2019 and the cross-cutting Inserm program on aging (AgeMed). SA’s Lab is supported by grants from the INSERM, Ministère de la Recherche, Agence Nationale pour la Recherche (ANR), Institut National du Cancer (INCA), Fondation pour la Recherche sur la Cancer (ARC) and Fondation pour la Recherche Médicale (FRM). ALV’s lab is supported by grants from ANR (Lustra), INCa (PLBIO2019), EDF (CT9818) and La Ligue contre le cancer-Paris (RS21/75-24). SCA is a recipient of a European CO-FUND PhD fellowship from Institut Curie. IF’s lab was funded by grants from the Spanish Ministry of Science and Innovation (PID2019-110339RB-I00) and the Comunidad de Madrid (S2017/BMD-3875). EG’s lab was supported by the Fondation ARC (Program ARC), the ANR grants TELOPOST and the cross-cutting Inserm program on aging (AgeMed). This work was performed using the PICMI facility of IRCAN (supported by FEDER, ARC, le Ministère de l’Enseignement Supérieur, la région Provence Alpes-Côte d’Azur and Inserm). CIPHE is supported by PHENOMIN (French National Infrastructure for mouse Phenogenomics; ANR10-INBS-07). The imaging studies carried out on the TrGET platform (IPC/CRCM) received financial support from ITMO Cancer as part of the 3rd Cancer Plan for the acquisition of dedicated equipment.

## Author contributions

Conceptualization, M.B., S.A., A.L.V., I.F. and V.G; Methodology, M.B., D.C., C.C., C.F., S.C.A., S.B., F.J., L.B., F.F., R.C., E.J., C.S.F., G.G.; Investigation and validation, M.B., D.C., C.C., C. F., C.S.F., G.G; I.F., E. G., A. L. V., S. A., V.G.; Supervision, M.B., I.F., A.L.V, S.A., V.G, Project administration, A.L.V., S.A., V.G.; Funding acquisition, I.F., E.G., A.L.V., S.A., V.G, Writing-original draft, VG; Writing Review-Editing, M.B., C.C., D.C., I.F., A.L.V., S.A.

## Methods

### Materials Availability

Mouse lines generated in this study are deposited at Ciphe, Marseille, France. Details and code numbers appear in the Key Ressources Table. This study did not generate unique reagents.

### Data and Code Availability

Original data for single cell RNA-sequencing is available at the Gene Expression Omnibus (GEO), GSE165218.

### Ethical statement

Mice were used according to institutional guidelines, which complied with national and international regulations. All animals received care according to institutional guidelines, and all experiments were approved by the Institutional Ethics committee number 16, Paris, France (licence number 16-093).

### Mice

Mice were bred and maintained in specific-pathogen-free conditions with sterilized food and water provided ad libitum and were maintained on a 12 h light and 12 h dark cycle in accordance with institutional guidelines. Male mice were used in this study, at the age of 4 (young group) and 18 (old group) month old. After genotyping, p21^+/+^ and p21^+/Tert^ mice from each litter were randomly assigned to the young or old group until the sufficient number of individuals per group was reached. P21^+/−^ mice generated at the same time in the same conditions followed the same procedure.

### Primary pulmonary artery smooth muscle cells

Pulmonary artery smooth muscle were extracted and cultured as already described ^61^. Cells were cultured at 37°C, 5% O2, in DMEM (High glucose, -Pyruvate, Gibco) supplemented with 10%(v/v) decomplemented FBS and 1%(v/v) Penicillin-Streptomycin.

### mCherry-2A-mTert vector and mice generation

A 6,4 kb genomic fragment encompassing coding exons 2 and 3 of the *Cdkn1a* gene was isolated from a BAC clone of B6 origin (clone n° RP23-412O16; http://www.lifesciences.sourcebioscience.com) and subcloned into the *Not* I site of pBluescript II. The genomic sequence containing the homologous arms was checked by DNA sequencing. A synthetic gene encoding the mCherry, 2A peptide, and the 5’ coding region of mTert (beyond the *Sac* II site) was constructed by gene synthesis (By DNA2.0 Inc) to produce the mCherry-2A-mTert^*Sac*II^ cassette. Using ET recombination, the synthetic mCherry-2A-mTert^*Sac*II^ cassette was introduced in the 5’ coding region of the *Cdkn1a* gene replacing the start codon. The fulllength mTert was next introduced in the mCherry-2A-mTert^*Sac*II^ truncated cassette by inserting into the targeting vector a synthetic mTert (*Sac* II-*EcoR* I) fragment restoring the full-length mTert to give rise to the complete mCherry-2A-mTert cassette. A self-deleter NeoR cassette was further introduced in the *EcoR* I site of the targeting vector immediately after the mTert sequence. A cassette coding for the diphtheria toxin fragment A expression cassette was finally introduced in the *Not* I site of the targeting vector. All the elements of the final targeting are shown in Fig. S1a.

JM8.F6 C57BL/6N ES cells were electroporated with the targeting vector linearized with Fse1 ^62^. The scheme of the introduction of the mCherry-2A-mTert-Neo^R^ cassette in place of the ATG codon of Cdkn1a is shown in Fig. S1b. After selection with G418, ES cell clones were screened for proper homologous recombination. The correct integration of the cassette was verified by Southern blot with a Cdkn1a probe (Fig. S1c) and by long range PCR (Fig. S1d). The primers used to check the correct insertion of the cassette were:

1310_ScES_01 Fwd: 5’ CTGAATGAACTGCAGGACGA ;

1310_ScES_01_Rev : 5’ CTTGCCTATATTGCTCAAGG.

To ensure that adventitious non-homologous recombination events had not occurred in the selected ES clones, a Southern blot was performed with a mCherry probe (Fig. S1e).

Properly recombined ES cells were injected into Balb/c blastocysts. Germline transmission led to the self-excision of the loxP-Cre-NeoR-loxP cassette in male germinal cells. p21-mCherry-2A-mTert mice (p21^+/Tert^) were identified by PCR of tail DNA. The primers used for genotyping were as follows: sense-WT 5’-GCTGAACTCAACACCCACCT-3’, sense-p21-Tert 5’-GGACCTCTGAGGACAGCCCAAA-3’ and antisense 5’-GCAGCAGGGCAGAGGAAGTA-3’. The resulting PCR products were 435 (WT) and 520 (p21^+/*Tert*^) base pairs long. The proteins produced by the mRNA mcherry-2A-mTert polycistronic mRNA are shown in Fig S1f In the final construct, Cdkn1a protein is no further produced from the mutated allele because of the substitution of the C*dkn1a* [ATG] codon by the cassette and the presence of a STOP codon at the end of Tert.

As control mice, we developed p21^+/+^ littermates and p21^+/−^ mice that were obtained by crossing the p21 homozygous knockout strain *B6.129S6(Cg)-Cdkn1a^tm1led^/J* (from JAX laboratories) with isogenic p21^+/+^ mice. In the p21^+/−^ mouse, a neo cassette replaces exon 2 of *Cdkn1a.* Homozygotes are viable, fertile, and of normal size.

p21^+/*TertCI*^ mice were constructed with exactly the same targeting vector used to construct p21^+/*Tert*^ except that the codon GAT in *Tert* coding for D702 has been changed into GCA coding for A. Primers to genotype p21^+/*Tert*^ and *p21^+/TertCI^* mice are indicated in Supplemental Information.

### Verification of the TERT activity

To check that TERT expressed from the mCherry-2A-mTert can produce active telomerase, the mCherry-2A-mTert was amplified by PCR (S 5’-atatatgaattcatggtgagcaagggcgaggaggata; AS 5’-atatatgcggccgcttagtccaaaatggtctgaaagtct) from the targeting vector and cloned into the *EcoR* I/*Not* I sites of pCAG-GFP. In the resulting vector, the mCherry-2A-mTert cassette is expressed under the control of the strong constitutive CAG promoter. The resulting vector was transfected in the mTert-/- mouse ES cells (provided by Lea Harrington, Montreal). Prior to transfection, the Neo^R^ cassette in the Tert-/- ES cells was disrupted by CRISPR/CAS9 since the Tert-/- ES cells and the electroporated plasmid shared the same antibiotic resistance. The presence of telomerase activity was analysed in the cell extracts using the Telomere Repeat Amplification Protocol (TRAP) (see below). We could detect robust telomerase activity in mouse ES mTert-/- transfected by the construct pCAG-mCherry-2A-mTert (Fig. S2a).

### Fluorescence imaging of mice and organs

To next demonstrate *in vivo* the expression of the KI *Cdkn1a* allele in response to p21 promoter activation, littermate p21^+/+^ and p21^+/Tert^ mice were exposed either to a whole body ionizing radiation or to doxorubicin which both activate the p53-p21 axis of the DNA damage response (DDR) ^63^.

For irradiation experiments, p21^+/+^ and p21^+/Tert^ mice were imaged (time zero) and then exposed to 1.5-Gy of total body irradiation (RX, RS2000 Irradiator, Radsources). Non-invasive whole-body fluorescence (FLI) was determined at the indicated times post-treatment. Imaging was performed using a Photon-IMAGER (Biospace Lab), and mice were anesthetized with 3 % vetflurane through an O2 flow modulator for 5 min and then the image was acquired (4-s exposure, excitation=545nm, background=495nm, emission filter cut off=615nm). Corresponding color-scale photographs and color fluorescent images were superimposed and analyzed with M3vision (Biospace Lab) image analysis software.

For Doxorubicin treatments, the p21^+/+^ and p21^+/Tert^ mice were injected i.p. with doxorubicin (DXR) (20 mg/kg). Doxorubicin has been shown to activate p21 promoter to high levels in the liver and kidneys ^17^. 24h after DXR injection, mice were then sacrificed and organs were rapidly removed and imaged within 15 min of sacrifice (4-s exposure). After that, proteins and RNA were extracted from kidneys and liver for immunoblot (p21) and RT-qPCR (Tert) analyses, respectively.

### PA-SMC culture

Pulmonary artery smooth muscle were extracted and cultured as already described ^61^. At each passage, cells were counted and 50,000 cells were put back in culture in a 25 cm2 flask and cultured in DMEM supplemented with FBS. Representative images of cells in culture were obtained using EVOS M5000 Imaging System (ThermoFisher) on p21^+/+^ and p21^+/Tert^ cells 24h after plating on a 24 well plate.

### Animal preparation and lung histological analysis

Two groups of mice from the three genotypes were prepared in order to sacrifice 4-month-old mice and 18-month-old mice at the same time. To reach the final mice number per group and per genotype, this experiment was reproduced three individual times, on three independent cohort. For p21^+/Tert-CI^ mice, an independent cohort of p21^+/Tert^ and p21^+/Tert-CI^ mice was prepared. After mice sacrifice three lobes of the right lung were quickly removed and immediately snap-frozen in liquid nitrogen then stored at -80°C for biological measurements. Genomic DNA from the right lung of each animal was used for TESLA assay. The last lobe of the right lung was fixed with 2% formaldehyde (Sigma) and 0.2% glutaraldehyde (Sigma) for 45 minutes. Then, lungs were washed with PBS and stained in a titrated pH 6 solution containing 40 mM citric acid, 150 mM NaCl, 2 mM MgCl_2_, 5 mM potassium ferrocyanide, and 1 mg/ml X-Gal. Stained lobes were then imbedded in paraffin and 5μm thick sections were cut. After counter coloration of the nucleus with neutral red, 10 fields per section were acquired at an overall magnification of 500.

The left lungs were fixed by intratracheal infusion of 4% paraformaldehyde aqueous solution (Sigma) at a trans-pleural pressure of 30 cmH_2_O. For morphometry studies, 5 μm–thick sagittal sections along the greatest axis of the left lung were cut in a systematic manner to allow immunostaining. Lung emphysema was measured using mean linear intercept methods on hematoxylin-eosin-safran (HES) coloration as described by Weibel and Cruz-Drive. Briefly 20 fields/animal light microscope fields at an overall magnification of 500, were overlapped with a 42-point, 21-line eyepiece according to a systematic sampling method from a random starting point. To correct area values for shrinkage associated with fixation and paraffin processing, we used a factor of 1.22, calculated during a previous study. On the same fields, bronchial thickness was measured using ImageJ software on bronchus with lumen diameter <130μm. Ashcroft score was assessed as already described ^26^ on HES sections.

### Immunohistochemistry and immunofluorescence

For immunohistochemistry, slides were deparaffinized using HistoChoice (Sigma) and rehydrated using increasing concentration of ethanol. After antigen retrieval (10min, 95°C in citrate buffer, pH6) and cellular permeabilization (0.1% Triton X-100 in PBS, 10min), slides were incubated for 60 minutes in 1% BSA and 5% goat serum in PBS and then incubated overnight with anti-p21 antibody (BD Pharmingen, mouse, #556431) and anti-p16 antibody (abbiotec, rabbit, #250804). We used the ABC Vectastain kit (Vector Labs, Burlingame, CA) to mark the primary antibodies according to the user’s guide. The staining substrate was diaminobenzidine (FastDAB, Sigma-Aldrich, St. Louis, MO), and the sections were counterstained with hemalun. After counterstaining sections were mounted with coverslips.

For immunofluorescence, slides were incubated with citrate buffer 10mM at pH6 for 20min at 95C. Slides were then incubated with 2.5% horse serum (Vector Laboratory, S-2000) for 30min at 37°C, and then incubated with anti-isolectin B4 alexafluor 594 antibody (I21413, ThermoFischer Scientific), 2μg/ml in 2.5% horse serum. Slides were mounted with DAPI and observed under the microscope at a 20x magnification. For all immunostaining, 10 fields per lung were used for quantification.

### BrdU intraperitoneal injection and analysis

A subgroup of mice received an intraperitoneal injection of 5-bromo-2’-deoxyuridine (ab142567, abcam) diluted in PBS at 25mg/ml. Mice received 100mg/kg BrdU 24h prior to sacrifice by cervical dislocation. Organ samples were collected and fixed in formalin solution, then embedded in paraffin. BrdU immunostaining was performed following provider instruction, using BrdU immunohistochemistry kit (ab125306, abcam).

### RNA extraction, RT-qPCR, and Western blots

Total RNA was extracted using the method of ^64^ with reagents included in the RNeasy kit (Qiagen). To exclude contamination with genomic DNA, the RNA was treated with DNase I directly on mini columns. Reverse transcription was performed using 0.5 to 2.0 μg of RNA, 50 ng/μL random hexamers, and 200 U of Superscript IV (Invitrogen) in the 20 μL reaction volume for 15 min at 55°C, followed by inactivation for 10 min at 80°C. The resulting cDNA was diluted 5-20 fold and analyzed by real-time qPCR using the SYBR Green master mix (Takara bio) and 400 nanomolar each of the following primers: mTert-2253S 5’-AGCCAAGGCCAAGTCCACAA and mTert-2399A 5’-AGAGATGCTCTGCTCGATGACA to target all *Tert* transcripts/isoforms; 2A-F2 5’-AGCAGGAGATGTTGAAGAAAACCC and mTert-5’R2 5’-GGCCACACCTCCCGGTATC to target *mCherry-2A-Tert* transcript from the KI *Cdkn1a* allele; mActb-90S 5’-ACACCCGCCACCAGTTCG and mActb-283A 5’-GGCCTCGTCACCCACATAGG to target β actin transcript. At least one primer within each pair was designed to anneal onto the exon-exon junction to avoid priming on genomic DNA, and the specificity of amplification was verified on agarose gel.

Immunoblots were carried out using standard procedures. The membranes were incubated with primary antibodies followed by incubation with the secondary HRP conjugates, and the signal was detected using an enhanced chemiluminescence detection system (GE Healthcare). The unsaturated images were acquired using ChemiDoc MP imaging system (Bio-Rad), and the signal densities were quantified using Image Studio Lite ver 5.2 (LI-COR). Antibodies used were rabbit monoclonal anti-p21 [EPR3993] (abcam) and mouse anti-b-actin (SigmaAldrich A1978) antibody.

### Telomere Repeat Amplification Protocol (TRAP)

To measure telomerase activity in cellular/tissue extracts TRAP was performed according to the original protocol ^65,66^. Briefly, the cells or tissues were extracted on ice using standard CHAPS buffer (60 μL per 10^6^ cells) supplemented with a cocktail of protease inhibitors (Roche) and 20 units of RNase inhibitor (Applied Biosystems) followed by centrifugation at 18000 x g for 30 min at 4°C. The protein concentration in the supernatant was quantified using Pierce™ 660nm Protein Assay and the aliquots equivalent to 800, 400, and 200 ng of protein were used to extend 50 ng of the telomerase substrate (TS oligo) in the 25 μL reaction volume. For gel-based detection of the TRAP product we used FAM-labeled TS and the standard ACX and NT primers (Kim and Wu, 1997) to amplify the telomerase extension products and the internal TSNT control, respectively. For the realtime qPCR-based TRAP we used unlabeled TS and the ACX primer alone. The qTRAP was performed using the SYBR Green PCR mix, and the amount of CHAPS extracts was optimized by serial dilutions.

### Telomere Shortest Length Assay (TeSLA)

TeSLA was performed according to the protocol described by ^21^. Briefly, 50 ng of undigested genomic DNA was ligated with an equimolar mixture (50 pM each) of the six TeSLA-T oligonucleotides containing seven nucleotides of telomeric C-rich repeats at the 3’ end and 22 nucleotides of the unique sequence at the 5’ end. After overnight ligation at 35°C, genomic DNA was digested with *Cvi*AII, *Bfa*I, *Nde*I, and *Mse*I, the restriction enzymes creating either AT or TA overhangs. Digested DNA was then treated with Shrimp Alkaline Phosphatase to remove 5’ phosphate from each DNA fragment to avoid their ligation to each other during the subsequent adapter ligation. Upon heat-inactivation of phosphatase, partially double-stranded AT and TA adapters were added (final concentration 1 μM each) and ligated to the dephosphorylated fragments of genomic DNA at 16°C overnight. Following ligation of the adapters, genomic DNA was diluted to 20 pg/μL, and 2-4 μL was used in a 25 μL PCR reaction to amplify terminal fragments using primers complementary to the unique sequences at the 5’ ends of the TeSLA-T oligonucleotides and the AT/TA adapters. FailSafe polymerase mix (Epicenter) with 1 × FailSafe buffer H was used to amplify G-rich telomeric sequences. Entire PCR reactions were then loaded onto the 0.85% agarose gel for separation of the amplified fragments. To visualize telomeric fragments, the DNA was transferred from the gel onto the nylon membrane by Southern blotting procedure and hybridized with the ^32^P-labeled (CCCTAA)_3_ probe. The sizes of the telomeric fragments were quantified using TeSLA Quant software ^21^.

### Quantification And Statistical Analysis

Basic statistics, tests for the differences between the means, and one-way analysis of variance (ANOVA) were performed with the GraphPad Prism 7 Software. ANOVA followed by Bonferroni multiple comparison test was used to compare the means of more than two independent groups. Pairwise correlation and multiple regression analyses were performed using Statistica 13.0 software package (StatSoft/Dell).

### Single cell RNAseq

#### Isolation of lung single cells from 18 month-old p21^+/+^, p21^+/−^ and p21^+/Tert^ mice

Mouse trachea is injected with 1,5ml dispase 50 U/ml (Corning #354235) followed by 0,5 ml agarose 1%. Lung is resected and minced in 3 ml DPBS 1x CaCl^2+^ and MgCl^2+^ (Gibco, 14040-091). Collagenase/dispase 100 mg/ml (Roche #11097113001) is added to the chopped tissue and placed on a rotator at 37 °C for 30 min. Enzymatic activity is inhibited by adding 5 ml of DPBS 1x (Gibco, 14190-094) containing 10% FBS and 1mM EDTA (Sigma #E7889). Cell suspension is filtered through 100 μm nylon cell strainer (Fisher Scientific #11893402), treated by DNase I (Sigma D4527) and filtered again through a 40 μm nylon cell strainer (Fisher Scientific #11873402). Red blood cells are removed by red blood cell lysis (Invitrogen, 00-4333-57) treatment for 90s at room temperature. Finally, lung cells are centrifuged at 150*g*, 4 °C for 6 min, resuspended in DPBS 1x containing 0,02% BSA (Pan Biotech #P06-13911000) and counted in a Malassez.

#### Preparation of single cell sequencing libraries

Single-cell 3’-RNA-Seq samples were prepared using single cell V2 reagent kit and loaded in the Chromium controller according to standard manufacturer protocol (10x Genomics, PN-120237) to capture 6.000 cells. Briefly, dissociated lung cells are encapsulated using microfluidic device. RNAs are captured on beads coated of oligos containing an oligo-dTTT, UMIs and a specific barcode. After reverse transcription, cDNAs are washed, PCR-amplified and washed again before analysis on a Bioanalyzer (Agilent) for quality control. Finally, libraries are prepared following standard Illumina protocol and sequenced on a NovaSeq sequencer (Illumina). Raw sequences are demultiplexed and reads are mapped onto the mm10 reference genome using the v2.3 Cell Ranger pipeline (10X Genomics) to generate a count matrix for each sample.

#### Quality control and Normalization of the single-cell data

The digital matrices were filtered from ambient contaminating RNA using the SoupX package (version 1.4.8) (preprint biorxiv:^67^). Further filtering was done to remove low-quality cells with low UMI counts, doublets and cells with relatively high mitochondrial DNA content. Outlier analysis was performed with perCellQCMetrics from the scatter package. An upper cutoff was manually determined for each sample based on a plot of gene count versus UMI count or % of mitochondrial genes, to have at least 1000 UMIs, number of transcripts ranging between 1000 and 30000 and at most 14% mitochondrial transcripts. The quality was consistent across samples, and differences in RNA and gene content could be ascribed to cell-type-specific effects. DGE matrices from all samples (p21^+/+^, p21^+/−^ and p21^+/Tert^) sequenced simultaneously were merged and subsequently normalized using the deconvolution normalization method in the *scran R* package in order to correct for differences in read depth and library size inside and between samples.

##### Clustering, cell type, and cycle annotation

Seurat v3.2.2 was used to perform dimensionality reduction, clustering, and visualization on the unique combined (with all samples) and normalized matrix. After scaling the data, dimensionality reduction was performed using PCA on the highly variable genes. Seurat’s *FindNeighbors* function was run to identify cluster markers with the following parameters: reduction = “pca” and dims = 1:10, followed by the *FindClusters* function with resolution = 1. The Seurat function *RunUMAP* was used to generate 2-dimensional umap projections using the top principal components detected in the dataset. To annotate cells, we mapped each cluster to the Mouse Cell Atlas ^31^ by using the *scMCA* function ^68^. Cluster identities were further verified according to gene markers found with the *FindMarkers* function from Seurat. Doublet cells were identified manually as expressing markers for different cell types and the doublet subgroups were removed from the matrix. In order to define endothelial (EC) and AT2 cells subclusters, EC and AT2 cell classes were isolated and subjected to a new clustering by the *FindNeighbors* function with new parameters: reduction = “pca” and dims (1:15 for EC and 1:10 for AT2 cells). Seurat *FindAllMarkers* function was applied to each EC subclusters and the markers were compared to the top 50 gene markers previously found in lung EC ^32^. We thus annotated EC subclusters depending on the enrichment of marker genes and identified five subtypes of EC (artery, capillary_1, capillary_2, vein and lymphatic). We annotated two functionally divergent lung EC capillary clusters according to ^33^. We thus reassign EC capillary_1 as “aCap” and EC capillary_2 as “gCap”. In order to assign cell cycle phase to each cell, we used the Cyclone method to our single cell RNA-Seq dataset.

### Differential gene expression analysis

To identify differentially expressed genes, we used the Mann-Whitney test adjusted by the Benjamini & Hochberg (BH) method. When comparing 18 month-old p21^+/+^, p21^+/−^ and p21^+/Ter***t***^ EC expressing Cdkn1a (normalized count >0) versus those not expressing Cdkn1a (normalized count =0), significant genes were defined according to their adjusted p-value (< 0.05) and by the fact that these genes were expressed in at least 10% of the cells. For 3D representation (Fig. 7c), genes expressed in EC with −LOG(p.value)>4 only in the p21^+/Tert^ but not in the p21^+/+^, p21^+/−^ were considered for subsequent analysis.

*String* analysis (https://string-db.org) was performed to investigate interactions between the significant genes.

#### Pearson correlation analysis

Pearson correlation analysis was carried using a matrix of normalized counts per cell for each selected gene expressed in gCap cells. We performed pairwise-comparisons between the selected genes in 225, 323 or 265 cells from p21^+/+^, p21^+/−^ and p21^+/Tert^ gCap cells, respectively.

