## Supplementary information for "Telomerase Prevents Emphysema in Old Mice by Sustaining Subpopulations of Endothelial and AT2 Cells"

**SUPPLEMENTARY FIGURE LEGENDS**

**Figure S1. Construction of p21$^{+/Tert}$ knockin mice (related to Fig. 1a)**

**(a)** Upper panel, map of the targeting vector used the introduction of the mCherry-2A-Tert cassette (see Methods). The targeting vector comprises the 5' homologous arm, the mcherry-2A-Tert cassette, the self-deleter neoR cassette, the 3' homologous arm and the DT-A expression cassette.

**(b)** Scheme of the introduction of the *mCherry-2A-Tert-Neo*$^R$ cassette in place of the ATG codon of *Cdkn1a.* Yellow triangles surrounding the *Neo*$^R$ are the loxP sites. The diphtheria-toxin A expression cassette (DTA) is used a negative selection marker (see Methods).

**(c)** Checking the 5'integration of the *mCherry-2A-Tert-Neo*$^R$ cassette in Cdkn1a.
Upper panel, schematic representation of Cdkn1a allele before (top) and after the integration of the *mCherry-2A-Tert-Neo*$^R$ cassette (bottom). The two first Cdkn1a exons (Ex1, Ex2) are shown in blue. Ex2 is the first coding Exon. The 5' homologous arm is shown in red (top) or in brown (bottom). The probe used to check the 5' integration cassette is shown in green. The *Drd* I restriction sites used to check the integration are indicated. Lower panel, selected ES cell genomic DNAs were digested with *Drd* I and analysed by Southern blot with the Cdkn1a probe indicated in the upper panel. The WT Cdkn1a allele is expected to give a single band of 8528 bp while the insertion of the cassette produces an additional band of 7620 bp. The names of the clones are indicated.

**(d)** Checking the 3'integration of the *mCherry-2A-Tert-Neo*$^R$ cassette in Cdkn1a.
Left panel, schematic representation of Cdkn1a allele after the integration of the *mCherry-2A-Tert-Neo*$^R$ cassette. The blue arrow represents the Cdkn1a Exon 2 without its ATG. The 3' homologous arm is represented in brown. The positions of the primers used to check the correct insertion of the cassette are indicated in green. 1310_ScES_01 Fwd: 5' CTGAATGAACTGCAGGACGA ; 1310_ScES_01_Rev : 5' cttgcctatattgctcaagg. Right panel, Long Range PCR was performed with the indicated Fwd and Rev primers. The reverse primer was outside the 3' homologous arm. The WT allele produces no amplification while the mutant allele gives a band of 3240 bp. The clones are the same as in (C).

**(e)** Checking the single integration of the *mCherry-2A-Tert-Neo*$^R$ cassette. Genomic DNAs of the indicated clones were digested with *EcoR* V and analysed by Southern blot with a mCherry probe. DNA from the WT does not hybridize with the probe while the cassette produces a band of 5,88 kbp recognized by the mCherry probe.

**(f)** Proteins produced by the mRNA mcherry-2A-Tert polycistronic mRNA. Successful skipping produced the indicated proteins.

**Figure S2. Validation of the p21-Tert mouse model (related to Fig. 1B and C)**

**(a)** Evidence for transcriptional activation of the KI allele (*mCherry-Tert*) *in vivo* in response to doxorubicin. p21$^{+/+}$ and p21$^{+/TERT}$ mice received intraperitoneal injection of doxorubicin (20 mg/Kg). 24 hours after treatment, mice were sacrificed and liver and kidneys were harvested. *Tert* RT-qPCR experiments were performed using RNA extracted from livers and kidneys from p21$^{+/+}$ and p21$^{+/Tert}$ mice treated or not with doxorubicin.

**(b)** Induction of the *mCherry-Tert* cassette expression *in vivo* by treatment with doxorubicin in liver and kidneys.

**(c)** p21 expression and mCherry fluorescence in the liver and kidneys after doxorubicin treatments. Livers and kidneys of $p21^{+/+}$ and $p21^{+/Tert}$ mice were harvested 24 hours after doxorubicin treatment. The level of p21 protein was evaluated by semi-quantitative immunoblotting (upper panel) and the organs (liver and kidney) were imaged for mCherry fluorescence (lower panel). The colous scale is indicated in the figure.

**Figure S3. TeSLA Southern blots used for quantification of the Very Short Telomeres in the lungs of the young and old mice (Related to Fig. 3C)**

(**a**) Number of short telomeres per genome in lungs from the $p21^{+/+}$ and $p21^{+/Tert}$ littermates (4 and 18-month-old mice). Genomic DNA was extracted from whole lungs and the length of short telomeres was analysed by TeSLA. The fraction of short telomeres for each mouse (n=6 for each group) was estimated from cumulative frequency distributions for the following thresholds: 2.4 kb (red squares), 1.2 kb (yellow squares), 0.6 kb (green triangles) and 0.4 kb (blue circles). No threshold allowed the detection of significant difference using ANOVA. The individual southern blots are shown in (**c**).

**(b)** Correlation between short telomeres per genome (<1.2kb) and the SA-b-Gal positive cells (in %) for each mouse. Data are individual values with linear regression curve.

(**c**) TeSLA was initiated with 50 ng of genomic DNA extracted from mouse lungs. In the final PCR step, 500 pg of ligation product was used per reaction, and 9 independent reactions were performed for each sample (each lane correspond to an independent PCR reaction) to achieve > 100 amplified telomeres for quantification. Note that young mice, regardless of their genotype, have less telomeres shorter than 1 kb compared to the old mice.

**Figure S4**. **Single-cell analysis of lungs from WT, $p21^{+/-}$, and $p21^{+/Tert}$ 18 month-old mice**

(**a**) Unsupervised Uniform Manifold Approximation and Projection for Dimension Reduction (UMAP) clustering of lung cells. Lung cell populations were identified in lung samples from WT ($p21^{+/+}$), $p21^{+/-}$, and $p21^{+/Tert}$ 18 month-old mice by using Mouse Cell Atlas (MCA) annotation.

(**b**) Distribution for immune and non-immune cells in lungs from mice with the indicated genotypes.

(**c**) Dot plots of Cdkn1a expression in the different lung cell-types. Cell populations identities are shown on the y-axis. The size of the dots represents the fraction of the cells expressing Cdkn1a. The color code represents the average gene expression in p21-positive cells.

**Figure S5. Representative markers use to annotate lung endothelial cell classes (Related to Figure 7)**

EC classes were annotated according to specific markers defined by Kalucka et al[32]. Representative markers are indicated for each EC classes.

**Figure S6. Individual expression of DEG within each EC classes (related to Fig. 7)**
Expression of the selected genes in individual cells of the 5 EC classes is represented in violin plots. Capillary class 2 (gCap) has been subdivided in cluster 1 and 2.

**Figure S7. Heatmap of DEG in between of AT2 cells from WT, p21$^{+/-}$, and p21$^{+/Tert}$ lungs of 18 month-old mice (Related to Fig. 8)**
Heatmap of differentially expressed genes (DEGs) in single p21$^{+/Tert}$ AT2 cells compared to WT and p21$^{+/-}$. Columns represent individual cells, grouped by cluster. Light red, green, and blue bars delineate cells belonging to cluster 1, 2 and 3, respectively. Genes are grouped as described for Figure 9. Groups 1-4 correspond to genes that are up-regulated in p21$^{+/Tert}$ while groups 5-8 represent down-regulated genes. The right panel represents boxplot of gene expression level mean per cluster/genotype/group.

### SUPPLEMENTARY METHODS

#### MOUSE MODEL USED IN THIS STUDY

| **Name** | **Genotype** | **Origin** | **International identification** |
|---|---|---|---|
| p21$^{+/Tert}$ | Heterozygote, p$^{21+/Tert}$ | Ciphe, Marseille | 1310_mCherry-2A-mTert |
| p21$^{+/+}$ | littermate controls of p21$^{+/Tert}$ | Ciphe, Marseille | 1310_mCherry-2A-mTert |
| p21$^{+/-}$ | heterozygote for p21 | Jackson Laboratory | B6.129S6(Cg)-*Cdkn1a*$^{tm1Led}$/J |
| p21$^{+/TertCI}$ | Heterozygote p21$^{+/TertCI}$ | Ciphe, Marseille | B6-Cdkn1atm2Ciphe |

| **Genotyping** | **5' to 3'** | **size (bp)** |
|---|---|---|
| **WT p21 F** | GCTGAACTCAACACCCACCT | 435 |
| **WT p21 R** | GCAGCAGGGCAGAGGAAGTA | |
| **mutant p21-mCherry-2A-Tert F** | GGACCTCTGAGGACAGCCCAAA | 503 |
| **mutant p21-mCherry-2A-Tert R** | GCAGCAGGGCAGAGGAAGTA | |
| **mutant p21-mCherry-2A-Tert-CI F** | GGACCTCTGAGGACAGCCCAAA | 520 |
| **mutant p21-mCherry-2A-Tert-CI R** | GCAGCAGGGCAGAGGAAGTA | |
| **B6.129S6(Cg)-*Cdkn1a*$^{tm1Led}$/J F** | GTTGTCCTCGCCCTCATCTA | 240 |
| **B6.129S6(Cg)-*Cdkn1a*$^{tm1Led}$/J R** | GCCTATGTTGGGAAACCAGA | |
| **B6.129S6(Cg)-*Cdkn1a*$^{tm1Led}$/J F** | GTTGTCCTCGCCCTCATCTA | 447 |
| **B6.129S6(Cg)-*Cdkn1a*$^{tm1Led}$/J R** | CTGTCCATCTGCACGAGACTA | |

#### REAGENTS and RESOURCES

| REAGENT or RESOURCE | SOURCE | IDENTIFIER |
|---|---|---|
| Antibodies | | |
| rabbit p16-INK4 | abbiotec | Cat#250804 |
| mouse p21 | BD Pharmingen | Cat#556431 |

| | | |
|---|---|---|
| mouse phospho-Histone H2A.X (Ser139) | Millipore | Cat#05-636 |
| isolectinB4 alexafluor 594 | ThermoFisher | Cat#I21413 |
| rabbit p21 | Abcam | Cat#EPR3993 |
| beta-actin (from hybridoma) | Sigma | Cat#A1978 |
| Goat Anti-Mouse Immunoglobulins/HRP | Dako | Cat#P0447 |
| Goat Anti-Rabbit Immunoglobulins/HRP | Dako | Cat#P0448 |
| Donkey polyclonal anti Mouse Alexa555 | ThermoFisher | Cat#A31570 |
| Alexa488-OO-(TTAGGG)3 | PANAGENE | |
| Chemicals, Peptides, and Recombinant Proteins | | |
| Doxorubicin hydrochloride | Sigma | Cat#D1515 |
| 5-bromo-2’-deoxyuridine | Abcam | Cat#ab142567 |
| Dispase Bacillus polymyxa-derived | Corning | Cat#354235 |
| Collagenase/ Dispase | Roche | Cat#11097113001 |
| red blood cell lysis buffer | Invitrogen | Cat#00-4333-57 |
| DNaseI, type II | Sigma | Cat#D4527 |
| Critical Commercial Assays | | |
| ABC Vectastain kit | Vector Labs | Cat#PK-6100 |
| BrdU immunohistochemistry kit | Abcam | Cat#ab125306 |
| RNeasy kit | Qiagen | Cat#74104 |
| SYBR Green master mix | Takara | Cat#RR420 |
| V2 reagent kit | 10x Genomics | Cat#PN-120237 |
| Deposited Data | | |
| Experimental Models: Cell Lines | | |
| mTert-/- mouse ES cells | Lea Harrington, Montreal | |
| Experimental Models: Organisms/Strains | | |
| Mouse: $p21^{+/Tert}$ : B6.p21mCherry-2A-Tert | Ciphe, Marseille | 1310_mCherry-2A-mTert |
| Mouse: $p21^{+/-}$: B6.129S6(Cg)-*Cdkn1a*$^{tm1Led}$/J | Jackson Laboratory | B6.129S6(Cg)-*Cdkn1a*$^{tm1Led}$/J |
| Mouse: $p21^{+/Tert-CI}$: B6.p21mCherry-2A-Tert(D702A) | Ciphe, Marseille | B6-Cdkn1atm2Ciphe |
| Oligonucleotides | | |
| Software and Algorithms | | |
| M3Vision | Biospace Lab | |
| EVOS M5000 Imaging System | ThermoFisher | |
| NovaSeq sequencer | Illumina | |
| v2.3 Cell Ranger pipeline | 10x Genomics | |
| SoupX package (version 1.4.8) | Cran | |
| perCellQCMetrics | Roche | |
| Seurat v3.2.2 | Sajita Lab | |
| Prism 7 | GraphPad | |
| Statistica 13.0 software package | StatSoft/Dell | |

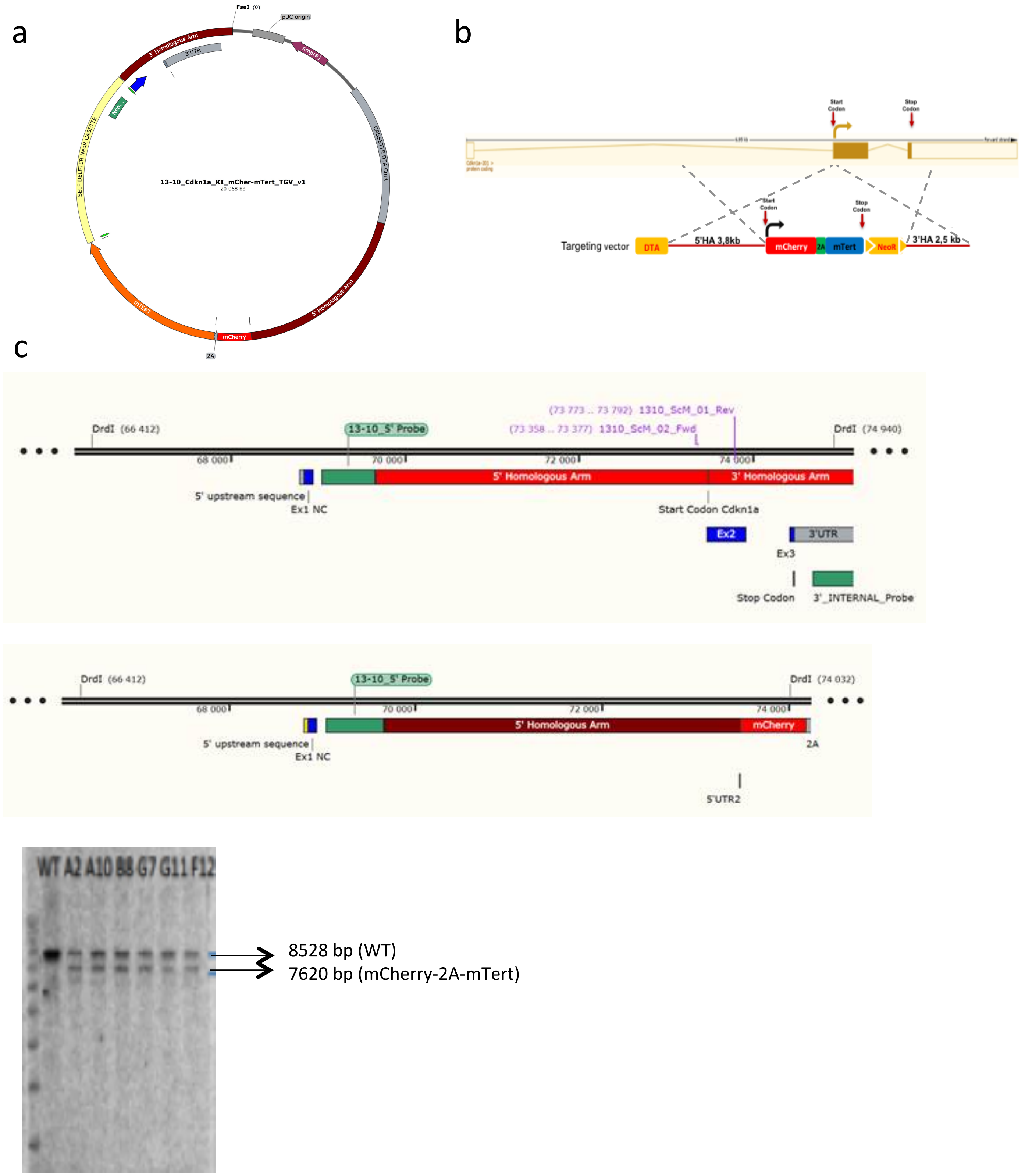

**Figure S1** (to be continued)

d

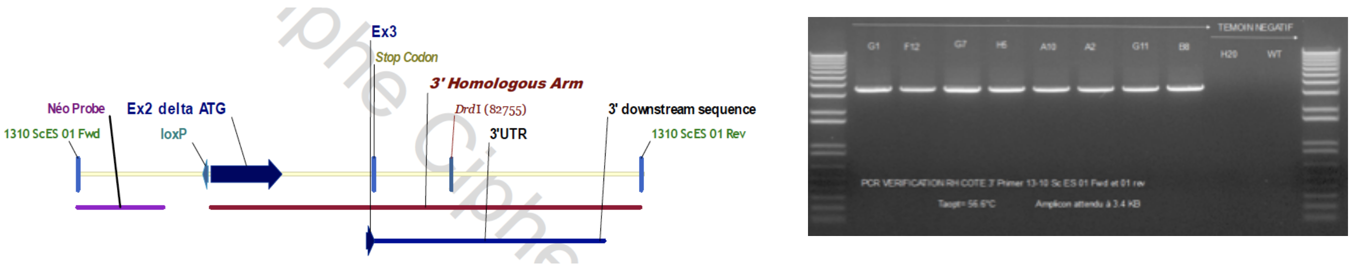

e

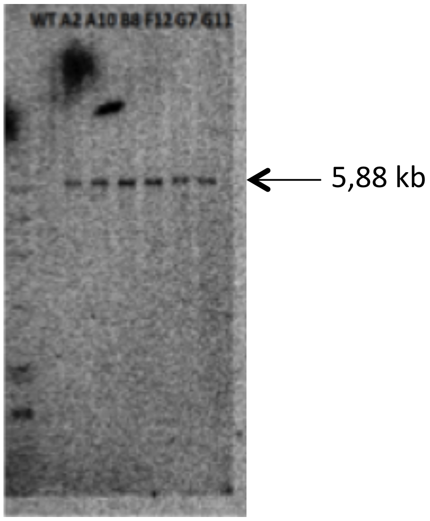

f

MVSKGEEDNMAIIKEFMRFKVHMEGSVNGHEFEIEGEGEGRPYEGTQTAKLKVTKGGPLPFAWDILS
PQFMYGSKAYVKHPADIPDYLKLSFPEGFKWERVMNFEDGGVVTVTQDSSLQDGEFIYKVKLRGTNF
PSDGPVMQKKTMGWEASSERMYPEDGALKGEIKQRLKLKDGGHYDAEVKTTYKAKKPVQLPGAYNVN
IKLDITSHNEDYTIVEQYERAEGRHSTGGMDELYKGATNFSLLKQAGDVEENPG

PLIKMTRAPRCPAVRSLLRSRYREVWPLATFVRRLGPEGRRLVQPGDPKIYRTLVAQCLVCMHWGSQ
PPPADLSFHQVSSLKELVARVVQRLCERNERNVLAFGFELLNEARGGPPMAFTSSVRSYLPNTVIET
LRVSGAWMLLLSRVGDDLLVYLLAHCALYLLVPPSCAYQVCGSPLYQICATTDIWPSVSASYRPTRP
VGRNFTNLRFLQQIKSSSRQEAPKPLALPSRGTKRHLSLTSTSVPSAKKARCYPVPRVEEGPHRQVL
PTPSGKSWVPSPARSPEVPTAEKDLSSKGKVSDLSLSGSVCCKHKPSSTSLLSPPRQNAFQLRPFIE
TRHFLYSRGDGQERLNPSFLLSNLQPNLTGARRLVEIIFLGSRPRTSGPLCRTHRLSRRYWQMRPLF
QQLLVNHAECQYVRLLRSHCRFRTANQQVTDALNTSPPHLMDLLRLHSSPWQVYGFLRACLCKVVSA
SLWGTRHNERRFFKNLKKFISLGKYGKLSLQELMWKMKVEDCHWLRSSPGKDRVPAAEHRLRERILA
TFLFWLMDTYVVQLLRSFFYITESTFQKNRLFFYRKSVWSKLQSIGVRQHLERVRLRELSQEEVRHH
QDTWLAMPICRLRFIPKPNGLRPIVNMSYSMGTRALGRRKQAQHFTQRLKTLFSMLNYERTKHPHLM
GSSVLGMNDIYRTWRAFVLRVRALDQTPRMYFVKADVTGAYDAIPQGKLVEVVANMIRHSESTYCIR
QYAVVRRDSQGQVHKSFRRQVTTLSDLQPYMGQFLKHLQDSDASALRNSVVIEQSISMNESSSSLFD
FFLHFLRHSVVKIGDRCYTQCQGIPQGSSLSTLLCSLCFGDMENKLFAEVQRDGLLLRFVDDFLLVT
PHLDQAKTFLSTLVHGVPEYGCMINLQKTVVNFPVEPGTLGGAAPYQLPAHCLFPWCGLLLDTQTLE
VFCDYSGYAQTSIKTSLTFQSVFKAGKTMRNKLLSVLRLKCHGLFLDLQVNSLQTVCINIYKIFLLQ
AYRFHACVIQLPFDQRVRKNLTFFLGIISSQASCCYAILKVKNPGMTLKASGSFPPEAAHWLCYQAF
LLKLAAHSVIYKCLLGPLRTAQKLLCRKLPEATMTILKAAADPALSTDFQTILD

*mCherry-2A*
*2A-Tert*

**Figure S1**

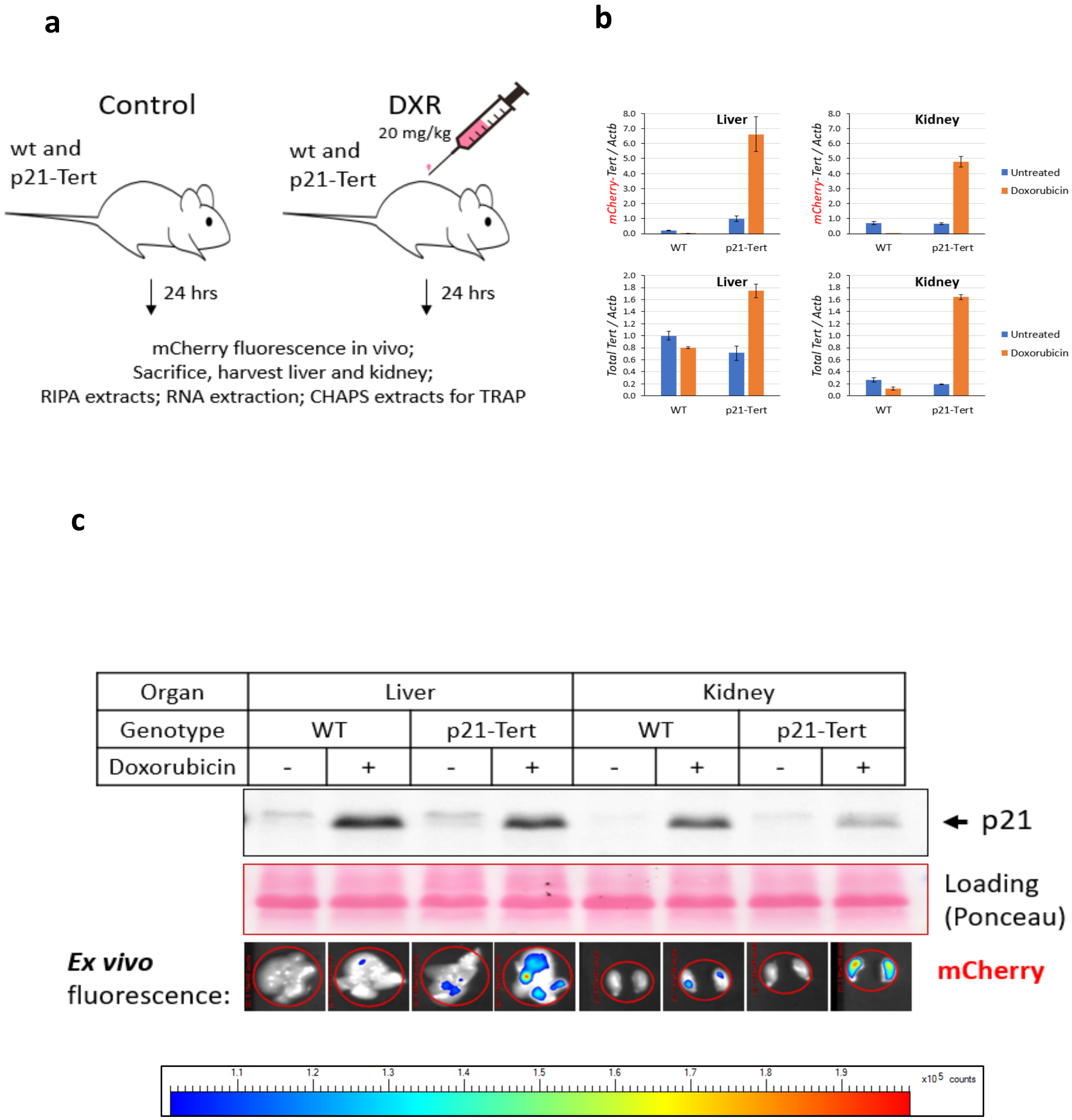
a
Control
wt and
p21-Tert
DXR
20 mg/kg
wt and
p21-Tert
24 hrs
24 hrs
mCherry fluorescence in vivo;
Sacrifice, harvest liver and kidney;
RIPA extracts; RNA extraction; CHAPS extracts for TRAP
b
Liver
Kidney
mCherry-Tert / Actb
Total Tert / Actb
WT
p21-Tert
Untreated
Doxorubicin
c
Organ
Liver
Kidney
Genotype
WT
p21-Tert
WT
p21-Tert
Doxorubicin
- + - + - + - +
p21
Loading
(Ponceau)
Ex vivo
fluorescence:
mCherry
1.1 1.2 1.3 1.4 1.5 1.6 1.7 1.8 1.9
x10 5 counts

Figure S2

a

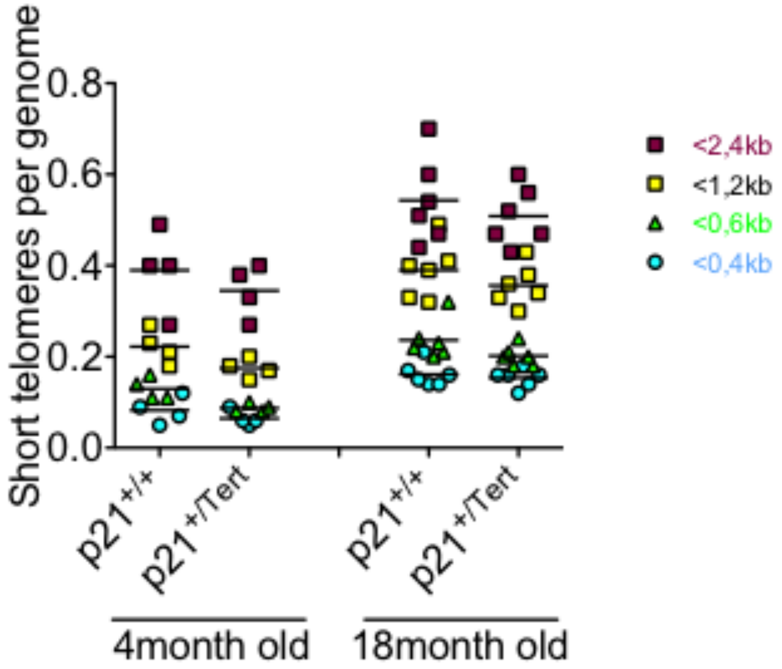

b

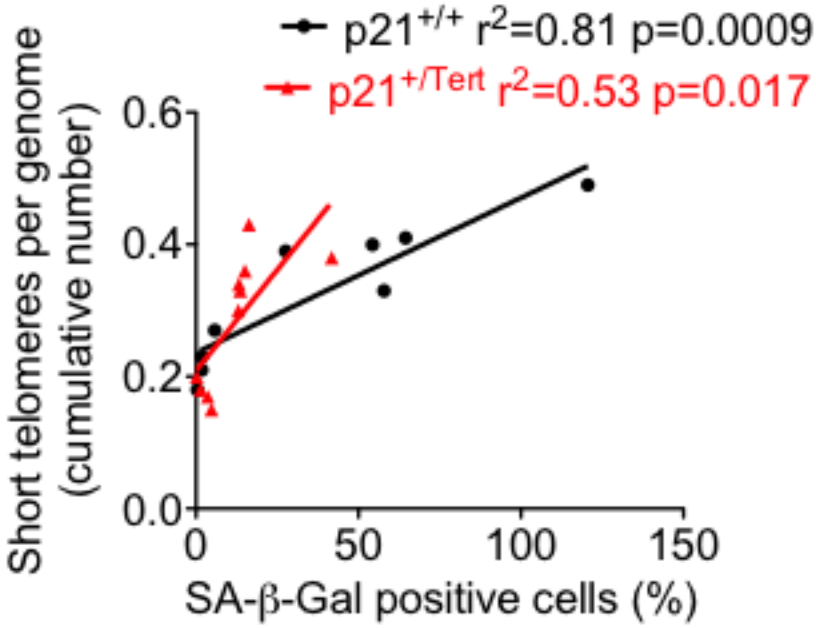

**Figure S3 (to be continued)**

c

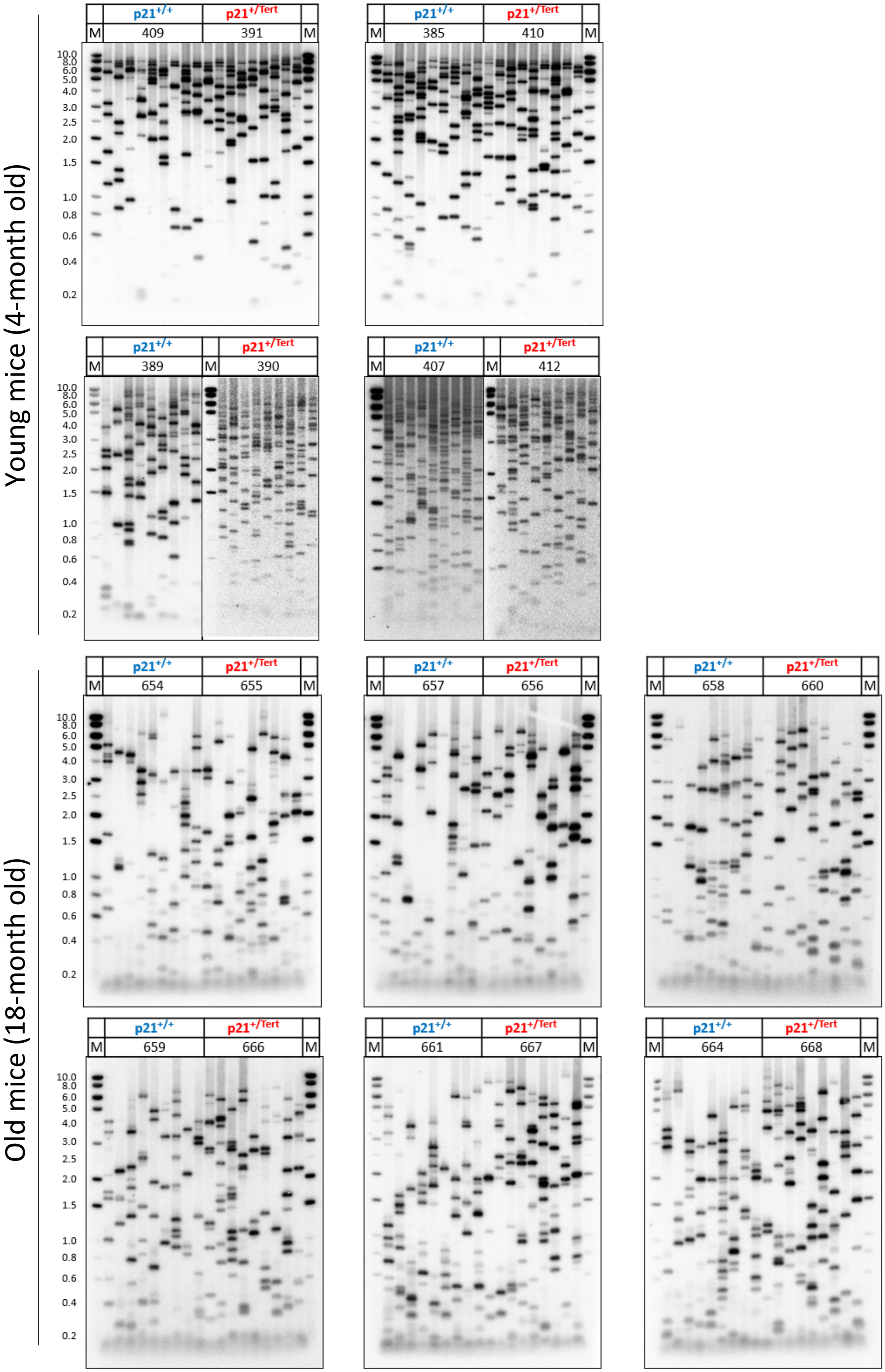

Young mice (4-month old)
Old mice (18-month old)
p21+/+
p21+/Tert
M
409
391
385
410
389
390
407
412
654
655
657
656
658
660
659
666
661
667
664
668
10.0
8.0
6.0
5.0
4.0
3.0
2.5
2.0
1.5
1.0
0.8
0.6
0.4
0.2

**Figure S3**

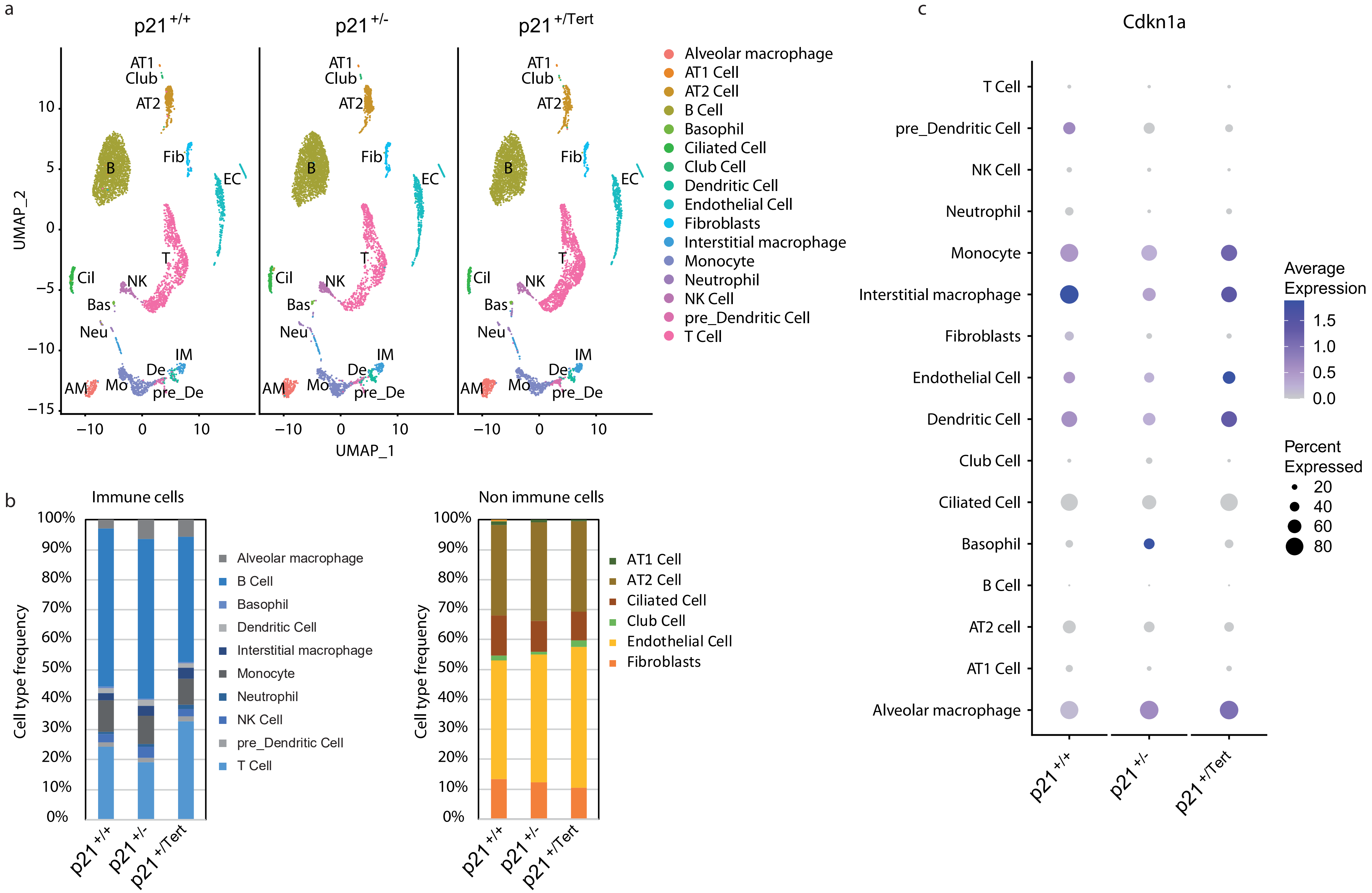
a
p21 +/+
p21 +/-
p21 +/Tert
UMAP_2
UMAP_1
AT1
Club
AT2
B
Fib
EC
T
Cil
NK
Bas
Neu
IM
De
AM
Mo
pre_De
Alveolar macrophage
AT1 Cell
AT2 Cell
B Cell
Basophil
Ciliated Cell
Club Cell
Dendritic Cell
Endothelial Cell
Fibroblasts
Interstitial macrophage
Monocyte
Neutrophil
NK Cell
pre_Dendritic Cell
T Cell
b
Immune cells
Cell type frequency
Alveolar macrophage
B Cell
Basophil
Dendritic Cell
Interstitial macrophage
Monocyte
Neutrophil
NK Cell
pre_Dendritic Cell
T Cell
Non immune cells
AT1 Cell
AT2 Cell
Ciliated Cell
Club Cell
Endothelial Cell
Fibroblasts
c
Cdkn1a
T Cell
pre_Dendritic Cell
NK Cell
Neutrophil
Monocyte
Interstitial macrophage
Fibroblasts
Endothelial Cell
Dendritic Cell
Club Cell
Ciliated Cell
Basophil
B Cell
AT2 cell
AT1 Cell
Alveolar macrophage
Average Expression
Percent Expressed

figure S4

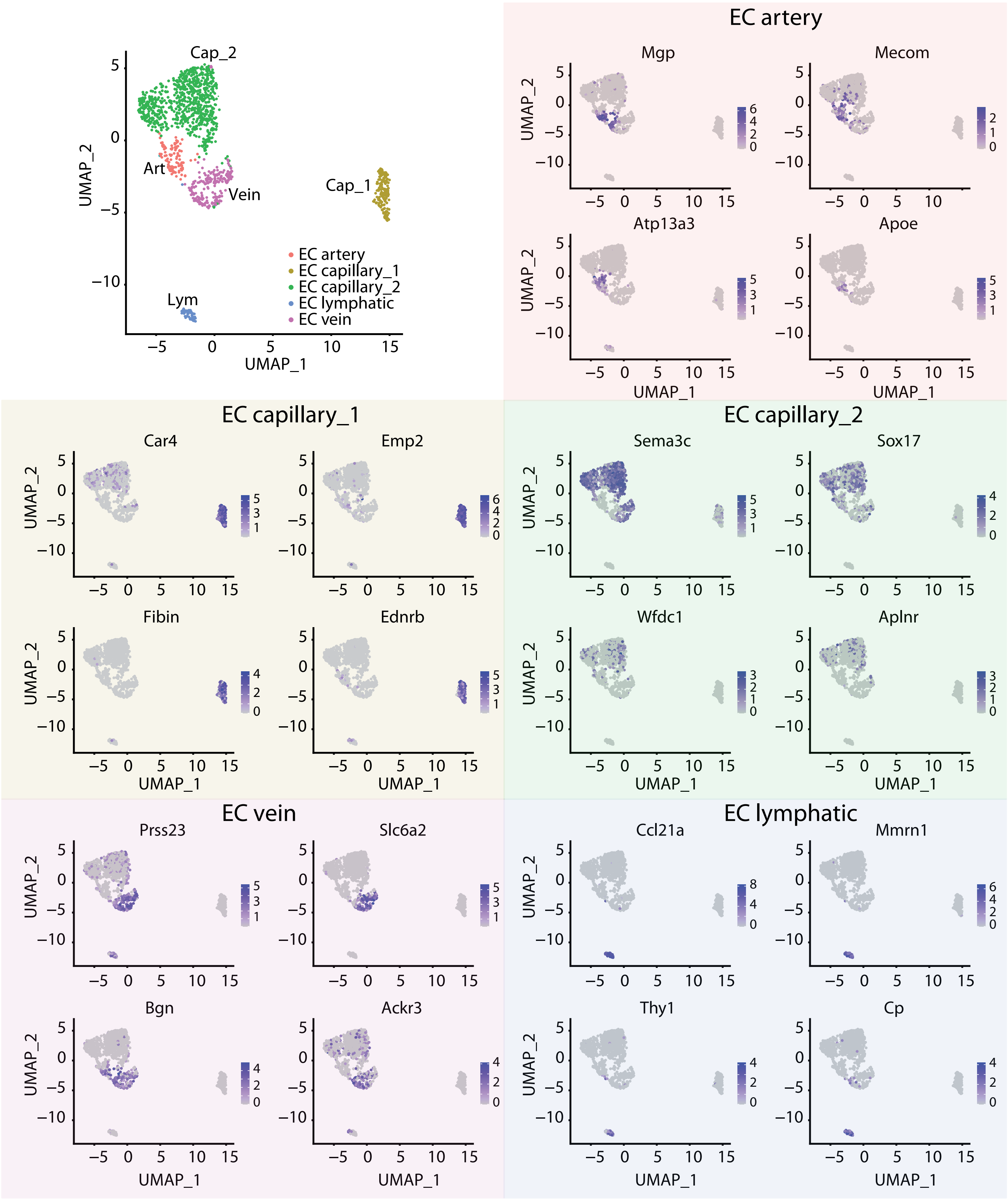
Cap_2
Art
Vein
Cap_1
Lym
EC artery
EC capillary_1
EC capillary_2
EC lymphatic
EC vein
UMAP_1
UMAP_2
EC artery
Mgp
Mecom
Atp13a3
Apoe
EC capillary_1
Car4
Emp2
Fibin
Ednrb
EC capillary_2
Sema3c
Sox17
Wfdc1
Aplnr
EC vein
Prss23
Slc6a2
Bgn
Ackr3
EC lymphatic
Ccl21a
Mmrn1
Thy1
Cp

figure S5

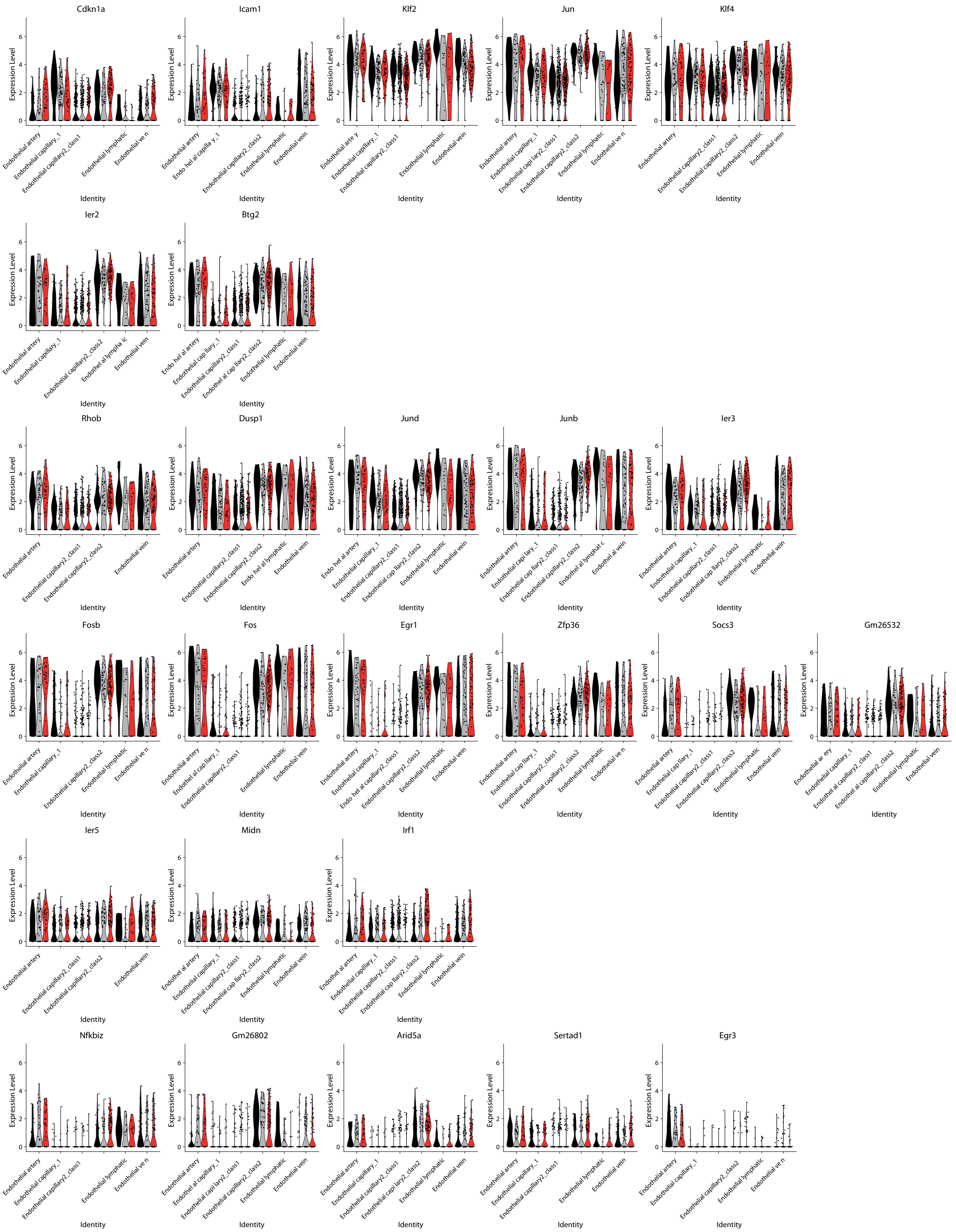
Cdkn1a
Icam1
Klf2
Jun
Klf4
Ier2
Btg2
Rhob
Dusp1
Jund
Junb
Ier3
Fosb
Fos
Egr1
Zfp36
Socs3
Gm26532
Ier5
Midn
Irf1
Nfkbiz
Gm26802
Arid5a
Sertad1
Egr3
Expression Level
Identity

figure S6

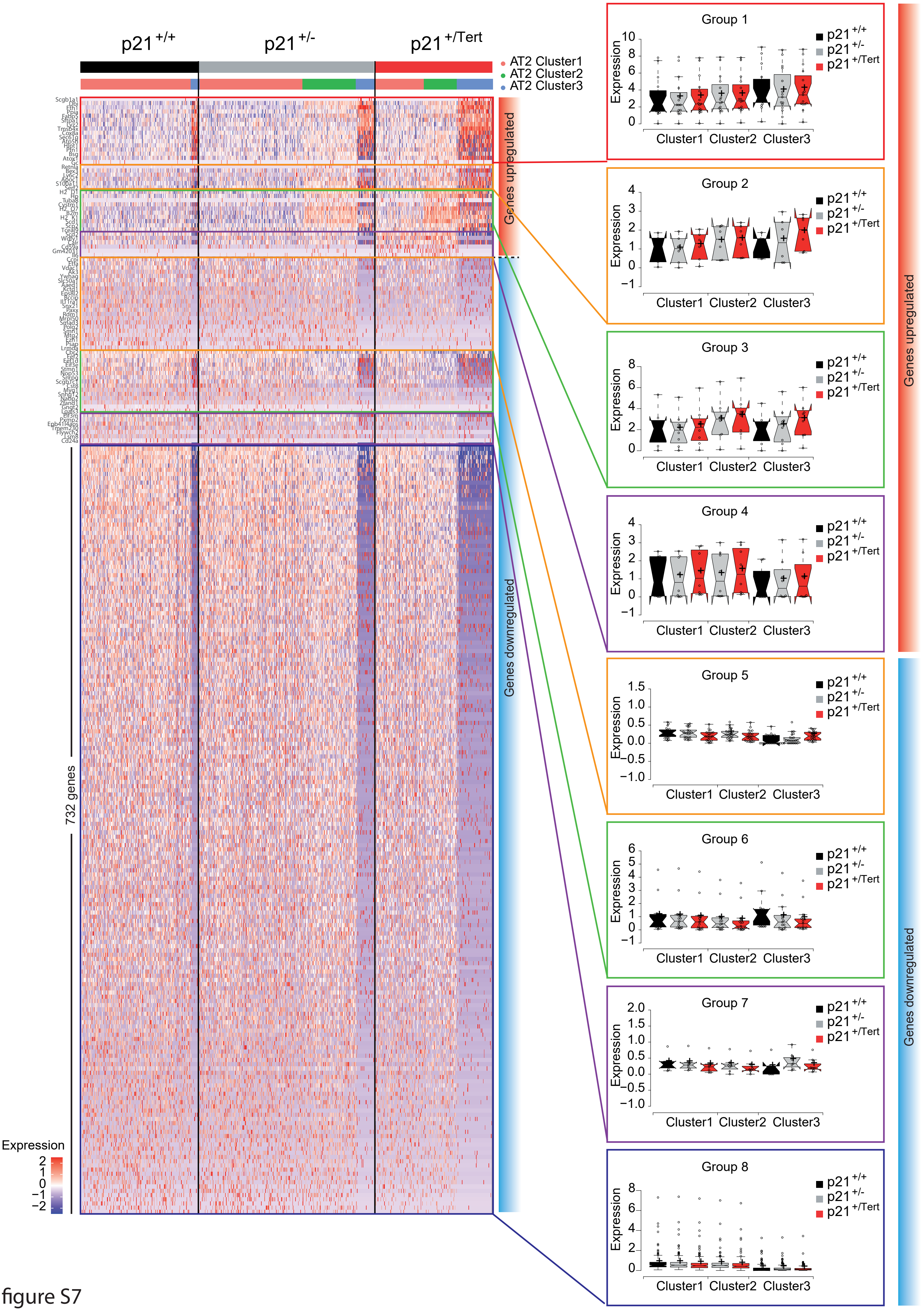

p21 +/+
p21 +/-
p21 +/Tert
AT2 Cluster1
AT2 Cluster2
AT2 Cluster3
Genes upregulated
Genes downregulated
732 genes
Expression
2
1
0
−1
−2
Group 1
Group 2
Group 3
Group 4
Group 5
Group 6
Group 7
Group 8
Expression
Cluster1 Cluster2 Cluster3
p21 +/+
p21 +/-
p21 +/Tert
Genes upregulated
Genes downregulated

figure S7
